# Bias-aware versus bias-blind confidence in humans and machines

**DOI:** 10.64898/2026.08.11.744086

**Authors:** Bogeng Song, Dobromir Rahnev

**Affiliations:** School of Psychological and Brain Sciences, Georgia Institute of Technology, Atlanta, GA

**Keywords:** Multi-alternative perceptual decision making, Confidence, Visual metacognition, Artificial neural networks, Bias monitoring

## Abstract

Confidence evaluates the likely accuracy of a current decision. However, to be maximally informative about accuracy, confidence judgments should incorporate information about one’s broader decision tendencies, such as their propensity to favor specific alternatives. We distinguish bias-aware confidence, which considers such tendencies, from bias-blind confidence, which relies only on evidence available on the current trial. To adjudicate between bias-aware and bias-blind confidence, we identified a signature of bias-aware confidence: the down-weighting of confidence for alternatives that a participant is biased toward. We then used a large dataset (N = 200) spanning 4- and 8-choice digit-classification tasks to show that humans reliably exhibit this signature of bias-aware confidence. This effect was reduced under speed pressure and could not be explained by guessing. In contrast to the human results, artificial neural networks (ANNs) trained for object recognition lacked this signature of bias- aware confidence. Importantly, augmenting ANNs with a metacognitive module that allows confidence to take the network’s biases into account led to the emergence of human-like bias- aware confidence. These findings show that human confidence incorporates not only information from the current trial but also longer-term decision tendencies, and that this capacity – absent in standard ANNs – can be conferred through specialized metacognitive mechanisms.

## Introduction

Perceptual confidence evaluates the likely accuracy of a specific decision (Fleming, 2024; Mamassian, 2016; Rahnev, 2021). However, decision accuracy depends not only on the current evidence strength but also on whether the available evidence reflects biases in sensory processing or decision-making. Such a bias can make the initial decision less reliable: if one alternative is frequently selected in error, then a trial on which that alternative is chosen is less likely to be correct. Therefore, to be maximally informative about accuracy, confidence should incorporate information about how one’s biases may have affected the available sensory evidence. We define confidence as “bias-aware” when it considers one’s own biases. Conversely, we define confidence as “bias-blind” when it relies only on evidence available on the current trial. Virtually all existing models of perceptual metacognition treat confidence as a readout of the strength of evidence on the current trial and thus tacitly assume bias-blind confidence (Hellmann et al., 2023; Maniscalco & Lau, 2016; Shekhar & Rahnev, 2024a). Yet, it remains unclear whether human confidence is indeed bias-blind or, instead, it incorporates mechanisms that account for one’s own biases.

What would bias-aware confidence look like in real-world decision making? Consider a radiologist interpreting an ambiguous opacity on a chest image. The opacity could reflect pneumonia, hemorrhage, malignancy, or atelectasis. Suppose the radiologist has the idiosyncratic tendency to diagnose pneumonia too readily. Bias-blind confidence would depend only on how strongly the current image seems to support pneumonia over the other diagnoses. In contrast, bias-aware confidence would also account for the radiologist’s tendency to over- select that diagnosis and would therefore discount confidence in it. Bias-aware confidence would thus allow radiologists to better evaluate the likely accuracy of their diagnosis, which could help them decide whether it would be worth running additional tests or consulting a colleague.

Several previous studies have examined confidence in the presence of response bias, but these studies induced bias through explicit reward or expectation cues (Constant et al., 2023, 2025; Lebreton et al., 2018; Locke et al., 2022; Schorn & Knowlton, 2026; Tarasi et al., 2025). One finding from this line of research is that external cues bias confidence judgments even more than the perceptual decisions. However, because in all these studies bias was induced externally through explicit cues, it remains unclear whether confidence reflects monitoring of internal bias or is simply sensitive to the external cues. Thus, it is critical to explicitly test whether human confidence is bias-aware in the sense of monitoring and taking into account one’s own natural biases.

Multi-alternative perceptual decisions provide a powerful way to adjudicate between bias- aware and bias-blind confidence. In two-choice tasks, bias toward one response is necessarily bias away from the other, making it difficult to isolate category-specific response tendencies. In contrast, multi-alternative tasks allow response bias to be estimated separately for each response category within each observer. This makes it possible to ask whether confidence is lower when a participant selects a response category that they tend to over-select. Critically, this approach tests whether confidence is sensitive to one’s own perceptual bias, rather than only to the current sensory evidence or to externally defined priors and rewards.

To directly adjudicate between bias-aware and bias-blind confidence, we first used multi- alternative signal detection theory (SDT) simulations to derive testable signatures of the two confidence strategies. The simulations showed that, after accounting for category-level accuracy, bias-blind confidence predicts a positive relationship between response bias and confidence, whereas bias-aware correction produces a negative relationship and improves the correspondence between confidence and accuracy. We then examined how humans give confidence in multi-alternative perceptual decision-making tasks. We collected a large dataset (N = 200) where participants performed 8- or 4-choice digit classification tasks using noisy images. We found that participants exhibited the signature of bias-aware confidence by down- weighting their confidence when they were biased toward a particular response. This effect was replicated in a separate dataset, which also revealed that speed pressure attenuates the relationship between response bias and confidence. Critically, standard artificial neural networks (ANNs) trained on object recognition instead exhibited bias-blind confidence. Finally, we demonstrated that bias-aware confidence emerges in ANNs enhanced with a metacognitive module that allows confidence to learn about the network’s response tendencies during training. These findings demonstrate that human confidence is bias-aware and that human-like confidence mechanisms can be incorporated into artificial systems through explicit metacognitive mechanisms.

## Results

### Basic confidence and response bias results

We tested N = 200 participants on a digit-classification task with two conditions that differed in the number of response alternatives: a 4-choice condition (digits 5–8) and an 8-choice condition (digits 1–8). On each trial, participants viewed a briefly presented noisy MNIST digit, indicated which digit they perceived, and then rated their confidence on a 1–4 scale (Figure 1A). Within each of the two conditions, the noise level in the images was adaptively adjusted using a 2-up-1-down staircase procedure to ensure a similar average accuracy across the two conditions. As intended, mean accuracy was closely matched between the two conditions (4- choice: 0.628 ± 0.013; 8-choice: 0.628 ± 0.015; t(199) = 0.49, p = 0.63). Further, we confirmed that confidence was significantly higher for correct than error trials (4-choice condition: *t*(199) = 42.41, *p* < .001; 8-choice condition: t(199) = 47.95, *p* < .001; Figure 1B), demonstrating that confidence ratings were meaningful.

**Figure 1.**
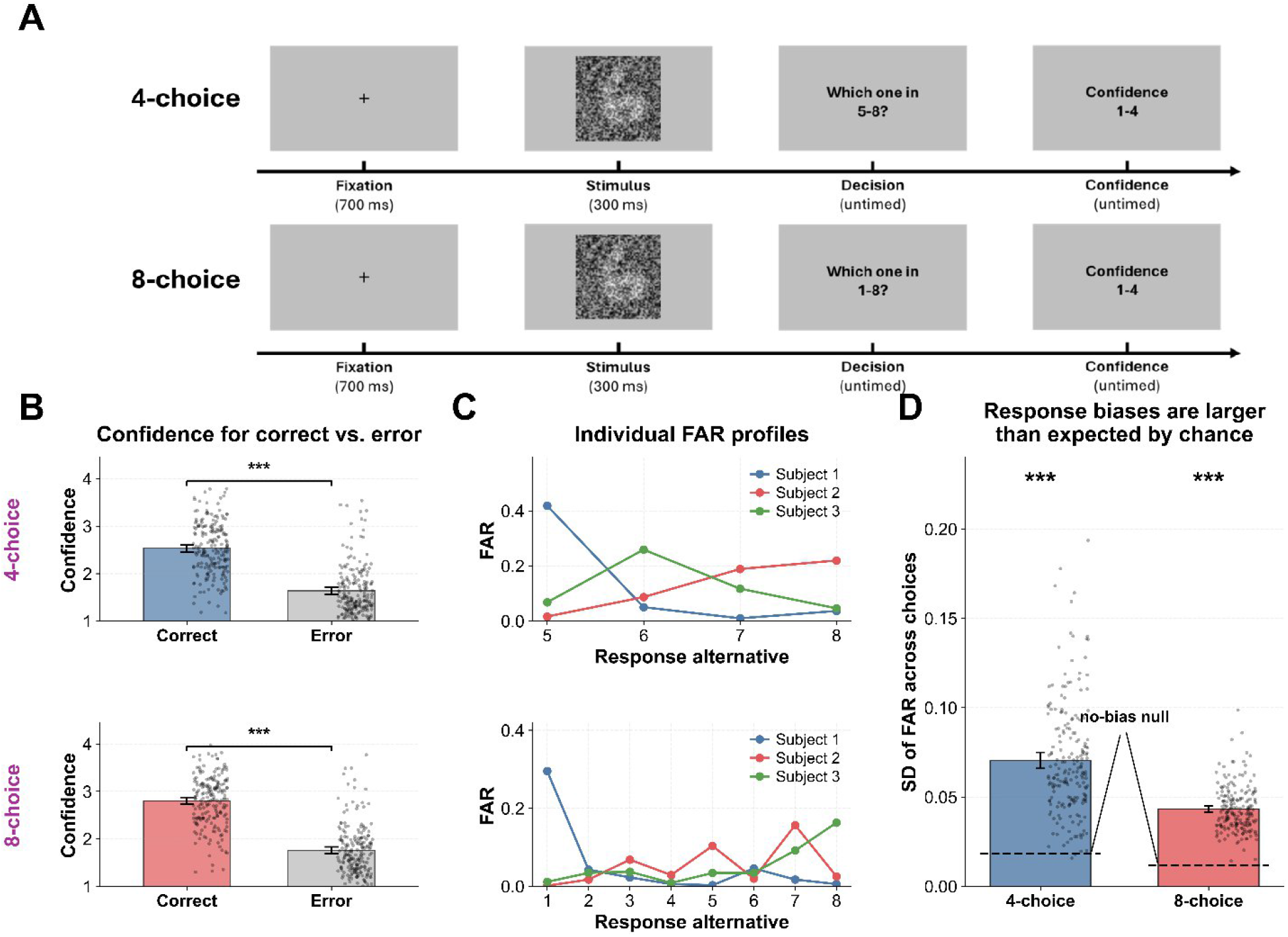
Task structure and basic behavioral results. (A) Task. Each trial began with a 700-ms fixation cross, followed by a 300-ms noisy digit stimulus. Participants then made an untimed digit-classification response and subsequently rated their confidence on a 1–4 scale. The top panel shows the 4-choice condition (digits 5–8), and the bottom panel shows the 8-choice condition (digits 1–8). (B) Confidence is higher for correct than error trials in both the 4- and 8- choice conditions. Points represent individual participants; error bars indicate SEM. (C) FAR profiles for three example participants in each condition demonstrating the presence of diverse biases. (D) Response biases are larger than expected by chance. The strength of response bias was quantified as the standard deviation of FAR across response alternatives. Bars show the observed mean across participants, dots represent individual participants, error bars indicate SEM, and dashed lines indicate the expected variability under uniform responding. ***, *p* < .001.

We then examined whether participants were biased to select some digits more than others. We computed response bias at the digit level as the false alarm rate (FAR) for that digit: that is, the proportion of trials on which that digit was selected, considering only trials on which a different digit was presented. Intuitively, a high FAR for a specific digit means that a participant is biased to select that digit even if it is not there. For illustration, we show the FAR patterns of three example participants in Figure 1C. To establish whether participants exhibited biases that are larger than expected by chance, we computed a null model by employing multi-alternative SDT simulations. We simulated data for 200 participants each completing 400 trials per condition, matching the human data. On a given trial, evidence for each incorrect category was sampled from *x_i_*_≠*p*_ ∼*N*(0,1) where *p* is the presented category, whereas evidence for the presented category was sampled from *x_p_* ∼*N* (*μ*,1) with the parameter *μ* chosen to produce accuracy that matches the average human accuracy. On each trial, the simulated participants simply selected the category with highest evidence, such that no response bias is introduced. We found that participants’ standard deviations across FAR values were substantially larger than predicted by the null model (4-choice condition: SD_empirical_ = .070, SD_null_ = .018; *t*(199) = 23.16, *p* < .001, Cohen’s d = 1.64; 8-choice condition: SD_empirical_ = .043, SD_null_ = .012; *t*(199) = 35.02, *p* < .001, Cohen’s d = 2.48; Figure 1D). In other words, the spread of FAR values exhibited by the participants was much higher than expected by chance, demonstrating that they were biased to over-select some digits and under-select others.

### A behavioral signature of bias-aware confidence

Having established the existence of substantial response bias in the empirical data, we turned to the question of whether human confidence judgments incorporate information about this bias. Conceptually, we would expect that such bias-aware confidence would feature down- weighting of confidence ratings for categories that an observer over-selects. We conducted simulations to examine how this conceptual effect translates into a quantitative signature that we can examine in the empirical data.

We employed the same multi-alternative SDT simulations as above but also added participant- specific response bias. Specifically, each simulated observer was also assigned a fixed, mean- centered response bias vector *b_i_* generated from a normal distribution *N* (0,*σ_b_*), where the value of *σ_b_* was selected to match the between-category spread of FAR in the empirical data. Thus, observers made their perceptual choices based on the biased evidence *e_i_* = *x_i_* + *b_i_*. The confidence variable included a correction for the estimated bias values 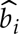 such that the confidence variable was 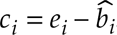. We simulated different levels of bias-aware correction such that 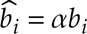 with the parameter *α* varying from 0 to 1 in steps of 0.1. Here, very low values of the parameter *α* correspond to bias-blind or mostly bias-blind confidence (i.e., no or minimal bias correction). In contrast, larger *α* values correspond to bias-aware confidence. Note that very high values of *α* are implausible as participants are unlikely to be able to perfectly estimate their own biases. The final confidence value was computed based on the difference between the chosen option and the highest among the remaining confidence evidence values (Li & Ma, 2020; Shekhar et al., 2025; Xue et al., 2024).

We found that without controlling for accuracy, the digit-level bias (FAR) always positively predicted the average digit-level confidence regardless of how strongly confidence corrected for bias (that is, regardless of the *α* value; Figure 2A). However, a similar regression where both digit-level FAR and digit-level accuracy jointly predicted average digit-level confidence showed a clear dissociation (Figure 2B). Specifically, low *α* values – corresponding to no or small confidence correction for bias – resulted in positive beta values for FAR, whereas high *α* values – corresponding to substantial confidence correction for bias – resulted in negative beta values for FAR. Further, the transition between positive and negative FAR beta values happened for *α* values between 0.5 and 0.6 for both the 4-choice and 8-choice conditions. These results demonstrate that the sign of the FAR beta value in a regression that includes both FAR and accuracy can be used as a signature of bias correction for confidence: negative beta coefficients demonstrate bias-aware confidence, whereas positive ones demonstrate mostly bias-blind confidence. Note that the simulations further uncovered an alternative measure of bias-aware confidence based on trial-level regression, which produced very similar results to the ones below (Supplementary Figures 1-3).

**Figure 2.**
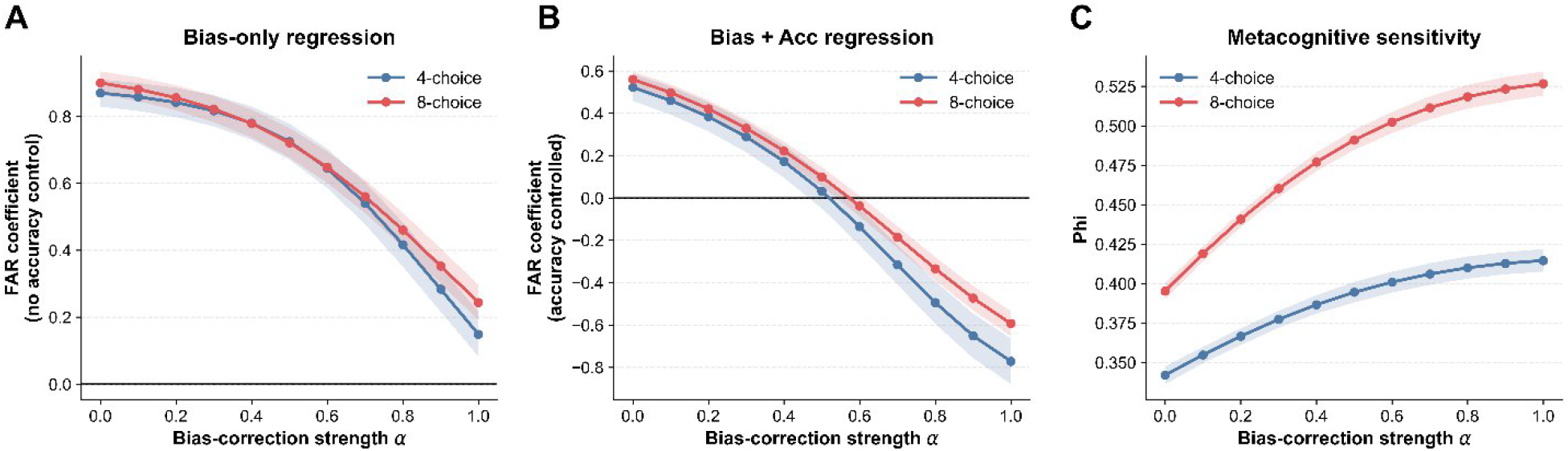
Multi-alternative SDT simulations identify a signature of bias-aware confidence. Simulated observers performed 4- or 8-choice perceptual decisions. We simulated the effects of different levels of bias-correction from none (*α* = 0) to perfect correction (*α* = 1). (A) FAR coefficients from mixed-effects regressions predicting category-level confidence from response bias alone, without controlling for accuracy. FAR positively predicted confidence across all *α* values. (B) FAR coefficients from mixed-effects regressions predicting category-level confidence from both FAR and category-level accuracy. FAR coefficients were positive at low levels of bias correction but became negative as *α* increased, with the sign reversal occurring between 0.5 and 0.6 in both choice conditions. (C) Metacognitive sensitivity, quantified as the mean within- observer correlation (Phi) between trial-level confidence and response correctness. Metacognitive sensitivity increased with bias-correction strength in both conditions. Shaded bands indicate 95% confidence intervals.

Importantly, our simulations allowed us to also examine whether correcting for bias makes confidence judgments align better with trial-by-trial accuracy. To address this question, we computed metacognitive sensitivity for different levels of correction. We quantified metacognitive sensitivity as Phi, the within-subject correlation between confidence and accuracy (Kornell et al., 2007; Rahnev, 2025). We found that stronger levels of bias correction led to higher metacognitive sensitivity (Figure 2C), demonstrating the advantages of confidence judgments that are correct for bias.

### Humans exhibit the signature of bias-aware confidence

Having established the predicted confidence signatures of bias-blind and bias-aware observers, we next examined whether humans exhibit the signature of bias-aware confidence. For illustration, we first consider the results for an example participant (Figure 3A). As expected, we found that digit-level bias, accuracy, and confidence are strongly correlated with each other. Critically, when accuracy was controlled for in a multiple regression as in the analysis in Figure 2B, we found the signature of bias-aware confidence: higher response bias became associated with lower confidence (4-choice: β = –0.97, 8-choice: β = –3.19; Figure 3A).

**Figure 3.**
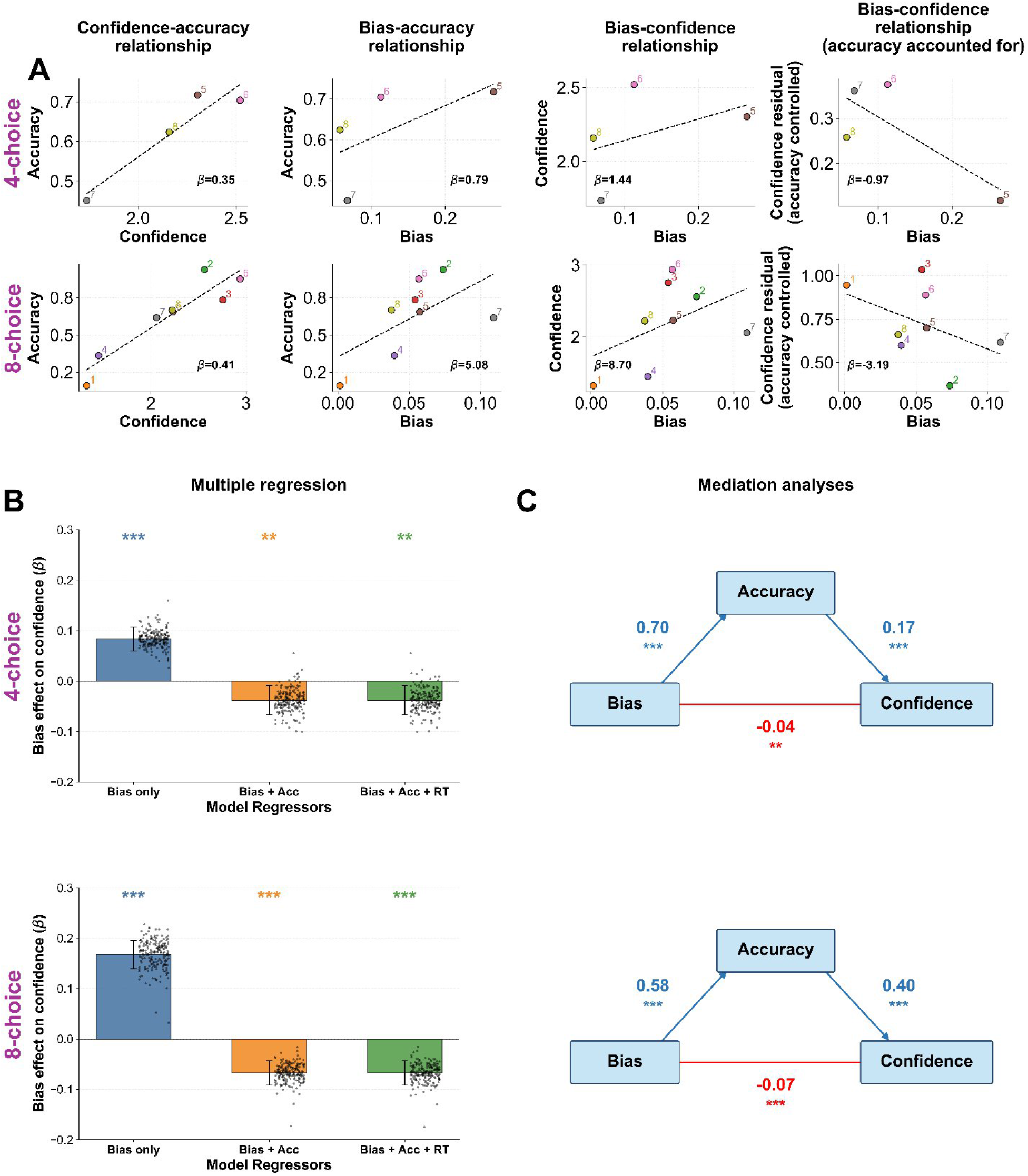
Response bias predicts lower confidence after accounting for accuracy. (A) Data from one representative participant in the 4-choice (top) and 8-choice (bottom) conditions. Bias was positively associated with both accuracy and confidence, but its relationship with confidence became negative after accuracy was controlled. Points represent digits, dashed lines show linear fits, and β values indicate regression slopes. (B) Mixed-effects regression estimates of the effect of bias on confidence. The bias effect was positive when bias was considered alone but became negative after controlling for accuracy and remained negative after additionally controlling for RT. Bars show mean β estimates, points show individual-participant estimates, and error bars indicate 95% confidence intervals. **, p < .01; ***, p < .001. (C) Bayesian mediation analyses show a negative direct effect of bias on confidence in both choice-set conditions when the indirect effect through accuracy is accounted for. Path coefficients represent posterior means. ** and *** indicate that the 99% and 99.9% highest density intervals, respectively, exclude zero.

We then turned to the population-level relationship between response bias and confidence. Replicating the pattern observed in the example participant, we found that higher response bias was associated with higher confidence when accuracy was not accounted for (4-choice: β = 0.083, SE = 0.012, z = 6.99, p < .001; 8-choice: β = 0.167, SE = 0.014, z = 11.81, p < .001; Figure 3B). However, once accuracy was controlled for in a multiple regression, the relationship between bias and confidence reversed direction, becoming significantly negative in both tasks (4-choice: β = –0.038, SE = 0.015, z = –2.58, p = .010; 8-choice: β = –0.067, SE = 0.012, z = –5.41, p < .001), thus exhibiting the signature of bias-aware confidence. Including response time as an additional covariate did not alter this pattern (4-choice: β = –0.038, SE = 0.015, z = –2.58, p = .010; 8-choice: β = –0.067, SE = 0.012, z = –5.42, p < .001).

To further examine the relationship between response bias, confidence, and accuracy, we performed a Bayesian mediation analysis (Figure 3C). Specifically, we examined the direct effect of response bias on confidence when the mediating effect of accuracy was accounted for. We found that accuracy has a strong mediating influence, such that there were strongly positive path coefficients from response bias to accuracy (4-choice: 0.16 [0.14, 0.19]; 8-choice: 0.39 [0.37, 0.41], mean and 95% HDI, Figure 3C) and from accuracy to confidence (4-choice: 0.69 [0.64, 0.74]; 8-choice: 0.57 [0.53, 0.61]). Critically, the direct effect of response bias onc onfidence was significantly negative (4-choice: –0.04 [–0.06, –0.01]; 8-choice: –0.07 [–0.09, –0.05]). Together, the multiple regression and mediation analyses show that response bias has a direct negative relationship with confidence, demonstrating the bias-aware nature of human confidence.

However, a possible alternative explanation of these results is that they are due to biased guessing accompanied by low confidence. Specifically, a strategy where participants always choose the same default digit when uncertain and report low confidence on such trials could also produce a negative relationship between response bias and confidence. Two additional analyses suggested that our results were not driven by such a strategy. First, we repeated the regression after removing trials with likely guesses based on RT. The logic was that putative guesses and lapses are likely associated with very fast or very slow responses (Ratcliff, 1993; Ratcliff & Tuerlinckx, 2002; Unsworth et al., 2010; Yellott, 1971). We performed a series of analyses by excluding all trials with RT under 300, 400, 500, 600, or 700 ms or RT above 2, 3, 4, or 5 seconds, and recomputed the FAR coefficient after controlling for accuracy and RT. Despite the reduced power of these analyses, we consistently observed a negative bias effect on confidence (Supplementary Figure 4A). Second, because guesses typically result in errors for multi-alternative decisions, we can examine confidence for correct trials only to remove the influence of putative guesses. Doing so also preserved the negative effect of bias on confidence (Supplementary Figure 4B). Together, these two control analyses strongly suggest that our results were driven by bias-aware confidence instead of low-confidence guessing.

### Speed focus attenuates the strength of bias monitoring in confidence

Bias-aware confidence requires that the current estimate of bias is retrieved and entered into the confidence computation. This process is likely to take time and resources. Therefore, the signature of bias-aware confidence – the negative effect of response bias on confidence after accounting for accuracy – should be reduced under conditions that restrict the time or resources available for the confidence computation.

To test this hypothesis, we analyzed the data from an existing dataset where N = 60 participants performed a similar 8-choice digit classification task. Critically, in different conditions participants were instructed to prioritize either accuracy or speed (Rafiei et al., 2024). As expected, participants exhibited faster response times (RTs) for the primary perceptual decision under speed compared to accuracy focus (mean RT = 1.06 vs. 0.86 s; t(59) = 11.16, p < .001, Cohen’s d = 1.44; Figure 4A). Crucially, we examined whether the speed condition also led participants to make faster confidence responses (S. Chen & Rahnev, 2023; Moran et al., 2015; Pleskac & Busemeyer, 2010). Indeed, confidence response times (cRTs) were shorter in the speed-focus compared to the accuracy-focus conditions (mean cRT = 0.51 vs. 0.56 s; t(59) = 2.07, p = .043, Cohen’s d = 0.27; Figure 4A). The shorter time for confidence responses would thus be expected to attenuate the extent of bias-correction exhibited by confidence judgments.

**Figure 4.**
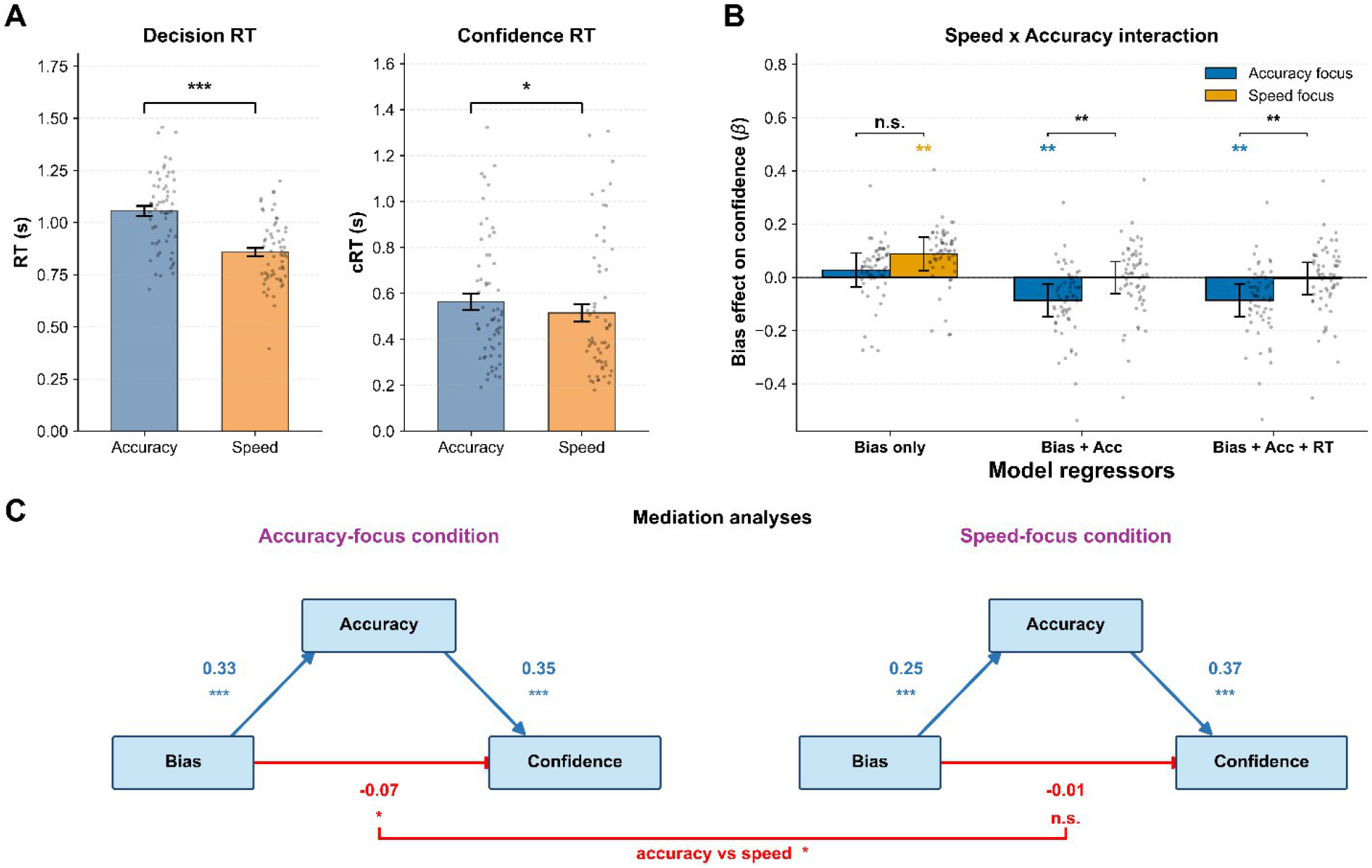
Speed focus attenuates the negative association between response bias and confidence. We re-analyzed the data from Rafiei et al. (2024), which included accuracy-focus and speed-focus conditions in an 8-choice digit classification task. (A) Mean decision RTs and confidence RTs (cRTs) under accuracy-focus and speed-focus instructions. Participants provided faster decisions and confidence judgments under speed focus than accuracy focus. Points represent individual participants, error bars show SEM. (B) Condition-specific response-bias slopes estimated from mixed-effects models fitted jointly to the speed-focus and accuracy- focus conditions. The negative effect of response bias on confidence when controlling for accuracy was attenuated under speed focus. This pattern remained after additionally accounting for RT. Points represent individual participants, error bars show 95% CIs. Asterisks above the bars indicate tests of condition-specific slopes against zero, whereas brackets indicate Bias × Condition interaction tests. (C) A joint Bayesian multilevel moderated mediation model showed that the direct path from response bias to confidence was negative under accuracy focus but near zero under speed focus. The bottom bracket indicates the posterior difference between the condition-specific direct effects. Path coefficients are posterior means. Panels A and B: *, p < .05; **, p < .01; ***, p < .001; n.s., not significant. Panel C: *, **, and *** indicate that the 95%, 99%, and 99.9% highest-density intervals excluded zero, respectively; n.s. indicates that the 95% highest-density interval included zero.

To test for such an effect, we fit both the speed-focus and accuracy-focus conditions in a single mixed-effects regression model with condition as moderator (Figure 4B). We found that condition indeed moderated the relationship between bias and confidence (β = –0.086, SE = 0.027, z = –3.24, p = .001). Specifically, in the accuracy-focus condition, response bias negatively predicted confidence after accounting for accuracy (β = –0.087, SE = 0.031, z = –2.76, p = .006), but this effect was not significant in the speed-focus condition (β = –0.001, SE = 0.031, z = –0.02, p = .983). This pattern remained virtually unchanged when RT was included in the model (Figure 4B). These results demonstrate that the signature of bias-aware confidence was indeed attenuated under speed pressure.

We further confirmed these findings using a joint Bayesian moderated mediation analysis (Figure 4C). We found that under accuracy focus, response bias had a negative direct effect on confidence (−0.07 [−0.12, −0.04]), whereas under speed focus, this direct effect was now indistinguishable from zero (−0.01 [−0.03, 0.05]). Critically, the moderation of the direct path was credibly negative (−0.09 [−0.14, −0.04]), confirming that the direct effect of response bias on confidence was different in the accuracy- vs. speed-focus conditions. Note that our simulations suggest that a direct effect of zero in our mediation analyses corresponds to a smaller bias correction instead of no bias correction (Supplementary Figure 5), meaning that the speed focus reduced rather than eliminated the bias-correction within confidence judgments. Together, the results of the regression and mediation analyses are consistent with the hypothesis that speed pressure reduces the ability of confidence ratings to take bias into account and thus attenuates the signature of bias-aware confidence.

### Bias-blind confidence in standard ANNs

Our results so far establish that humans exhibit bias-aware confidence. As a next step, we examined the bias-confidence relationship in a system that is a priori expected to exhibit bias- blind confidence: standard artificial neural networks (ANNs) trained on object recognition. Indeed, unlike humans whose confidence may incorporate information accumulated across previous decisions, standard ANNs derive confidence directly from the current trial’s output evidence (e.g., the relative magnitude of final-layer logits). Thus, these networks provide a computational model of bias-blind confidence, where confidence reflects current-trial evidence but does not adjust for systematic response tendencies associated with particular categories.

We trained three ANN architectures – AlexNet (Krizhevsky et al., 2012), ResNet18 (He et al., 2015), VGG19 (Simonyan & Zisserman, 2015) – on noiseless MNIST digit classification. We then examined how the response biases exhibited by these ANNs predicted their own confidence levels when tested on noisy images selected to produce accuracy levels comparable to that of our human participants. To establish robustness, for each architecture, we trained and tested 60 ANN instances that differed in their random weight initialization (Fung et al., 2025; Rafiei et al., 2024). Confidence was derived using the Top2Diff computation that considers the difference of the top two logits, as that computation has been suggested to most closely resemble the human confidence computation (Shekhar et al., 2025).

We found that all three ANNs exhibited the signature of bias-blind confidence: a positive direct effect of response bias on confidence (Figure 5). As expected, mixed-effects regression analyses showed that response bias positively predicted confidence when accuracy was not accounted for (AlexNet: β = 0.44, SE = 0.04, z = 11.75, p < .001; ResNet18: β = 0.44, SE = 0.04, z = 11.83, p < .001; VGG19: β = 0.70, SE = 0.08, z = 8.37, p < .001; Figure 5A). Critically, unlike for human participants, this relationship remained significantly positive even after controlling for accuracy (AlexNet: β = 0.32, SE = 0.04, z = 7.24, p < .001; ResNet18: β = 0.30, SE = 0.04, z = 6.76, p < .001; VGG19: β = 0.19, SE = 0.04, z = 4.45, p < .001). The same pattern emerged using our Bayesian mediation analyses (Figure 5B). Specifically, similar to humans, we found positive path coefficients from response bias to accuracy (AlexNet: 0.43 [0.37, 0.50]; ResNet18: 0.56 [0.47, 0.65]; VGG19: 0.82 [0.69, 0.94]) and from accuracy to confidence (AlexNet: 0.80 [0.55, 1.03]; ResNet18: 0.48 [0.29, 0.66]; VGG19: 0.92 [0.78, 1.08]). Crucially, and unlike humans, the direct effects of response bias on confidence were strongly positive across all ANNs (AlexNet: 0.36 [0.23, 0.49]; ResNet18: 0.44 [0.36, 0.53]; VGG19: 0.24 [0.14, 0.33]). These results were replicated using an alternative, trial-based regression analysis (Supplementary Figure 3), as well as when employing SoftMax instead of Top2Diff to derive confidence (Supplementary Figure 6). Overall, these findings show that standard ANNs exhibit bias-blind confidence: response bias exhibits a strong positive relationship to confidence even after accuracy is controlled.

**Figure 5.**
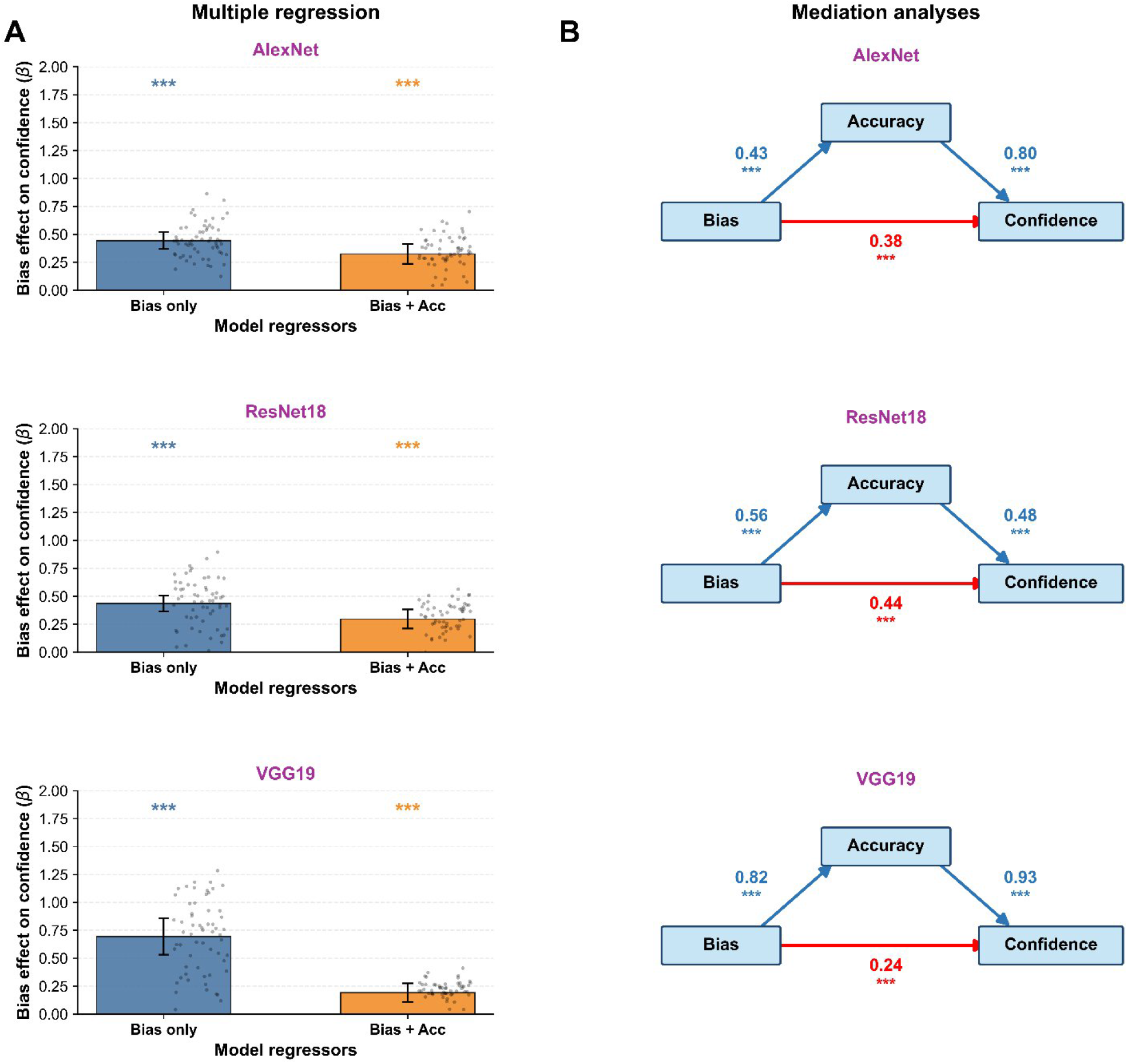
Standard ANNs show a positive effect of bias on confidence. Unlike humans who exhibit negative direct effect of response bias on confidence, this effect is positive in standard ANNs. (A) Mixed-effects regression for AlexNet, ResNet18, and VGG19. Response bias positively predicted confidence both before and after controlling for accuracy across all architectures. ***, p < .001. (B) Bayesian mediation analyses for AlexNet, ResNet18, and VGG19. Mediation analyses revealed robust positive direct effects of response bias to confidence. Path coefficients represent posterior means; asterisks indicate the 99.9% (***) highest-density intervals exclude zero.

### Adding a metacognitive module in ANNs produces bias-aware confidence

Given that standard ANNs exhibit bias-blind confidence, we explored whether we could augment ANNs with a metacognitive module to allow them to give confidence that corrects for their biases. For each architecture, we froze the trained base classifier and trained only an auxiliary metacognitive module to predict whether the classifier’s response to a noisy image was correct. For each image, the module was given the classifier’s final-layer logits and was trained to predict whether the classifier’s chosen response was correct. At test, the trained module produced a scalar confidence estimate for each trial. This approach follows the logic of learned confidence and failure-prediction models (T. Chen et al., 2019; Corbière et al., 2019), while preserving the base model’s perceptual processing and response policy. The metacognitive module could thus learn the classifier’s biases and account for them when producing confidence without affecting the underlying perceptual representations or response choices.

We found that this simple metacognitive module allowed ANNs to exhibit the signature of bias- aware confidence (Figure 6). First, as expected, when accuracy was not accounted for, response bias positively predicted confidence for all three architectures (AlexNet: β = 0.37, SE = 0.09, z = 3.92, p < .001; ResNet18: β = 0.54, SE = 0.07, z = 8.14, p < .001; VGG19: β = 0.82, SE = 0.07, z =11.97, p < .001). Critically, once accuracy was accounted for in the multiple regression, response bias showed a negative influence on confidence for all three architectures (AlexNet: β = –0.10, SE = 0.01, z = –14.76, p < .001; ResNet18: β = –0.05, SE = 0.004, z = –13.25, p < .001; VGG19: β = –0.05, SE = 0.005, z = –10.30, p < .001). The Bayesian mediation further confirmed the negative direct effect of response bias on confidence (AlexNet: –0.10 [–0.11, –0.09]; ResNet18: –0.05 [–0.06, –0.04]; VGG19: –0.05 [–0.06, –0.04]), while positive path coefficients remained from response bias to accuracy (AlexNet: 0.42 [0.36, 0.50]; ResNet18: 0.52 [0.44, 0.60]; VGG19: 0.80 [0.680, 0.94]) and from accuracy to confidence (AlexNet: 1.01 [1.00, 1.02]; ResNet18: 1.01 [1.00, 1.01]; VGG19: 1.00 [1.00, 1.01]). Therefore, adding a learned metacognitive readout to otherwise fixed ANNs led these networks to exhibit the signature of bias-aware confidence.

**Figure 6.**
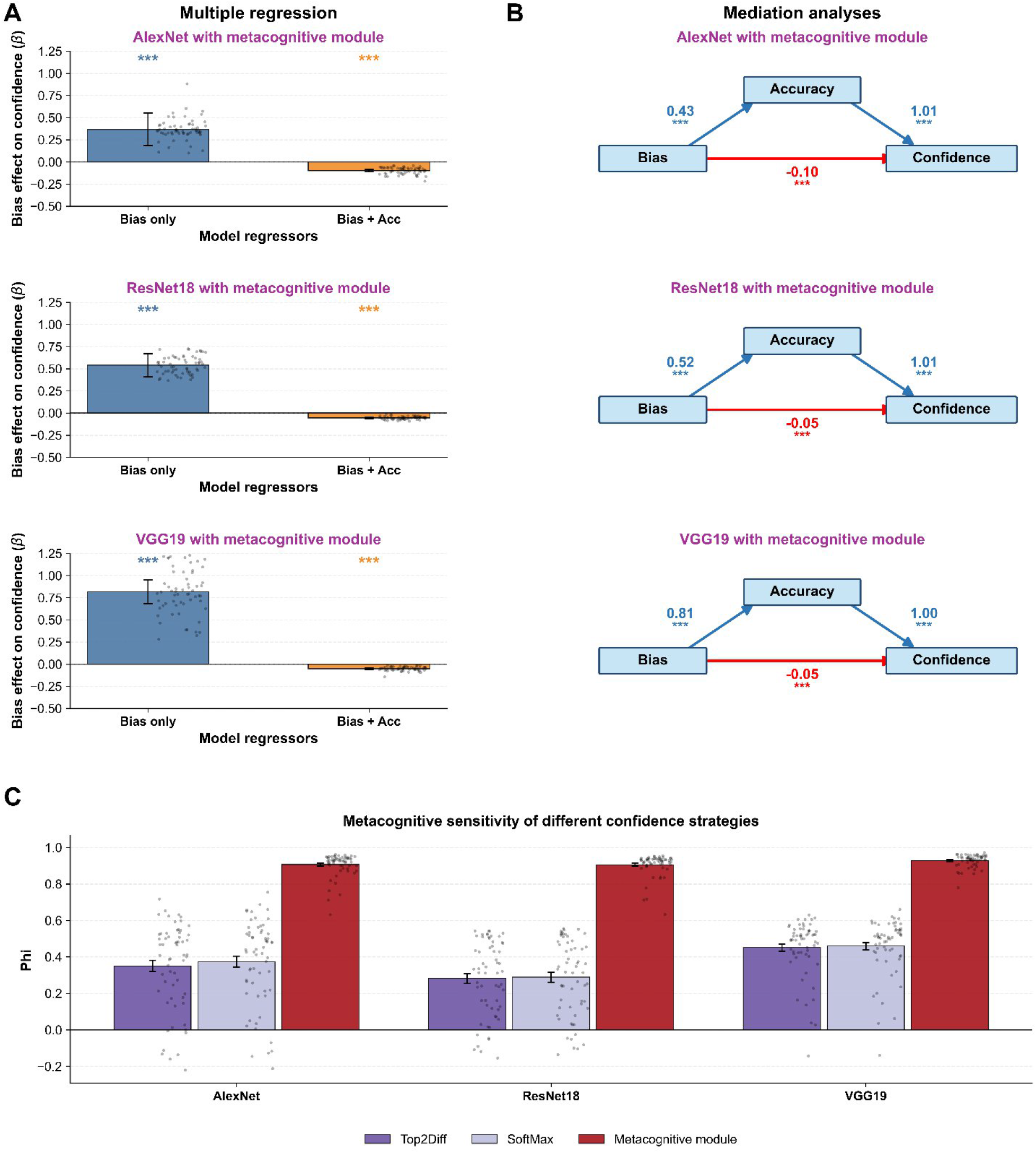
A metacognitive module produces human-like bias-aware signatures in ANNs. (A) Mixed-effects regressions predicting confidence from response bias for AlexNet, ResNet18, and VGG19 augmented with a metacognitive module. In the bias-only models, response bias positively predicted confidence in all three architectures. After accounting for accuracy, response bias negatively predicted confidence across architectures, reproducing the bias-aware confidence signature observed in humans. Points represent individual ANN instances; error bars show 95% confidence intervals; ***, p < .001. (B) Bayesian multilevel mediation analyses for the three architectures. The direct paths from response bias to confidence were negative across architectures, indicating that the metacognitive module adjusted confidence for the networks’ response tendencies. Path coefficients are posterior means; *** indicates that the 99.9% highest-density interval excluded zero. (C) Metacognitive sensitivity (Phi) for three confidence readouts. Phi was substantially greater when confidence was derived from the metacognitive module compared to the Top2Diff and SoftMax computations. Points represent individual ANN instances; error bars show SEM.

Finally, we examined whether the metacognitive module increased the ANNs’ metacognitive sensitivity, as would be expected for bias-aware confidence (Figure 2). We computed the confidence-accuracy correlation (Phi) for the ANNs using the Top2Diff and SoftMax computations (that do not include training a metacognitive module) and for the ANNs that included our metacognitive module. We found the metacognitive module produced substantially higher metacognitive sensitivity across all architectures (average Phi correlation across ANN architectures of 0.91, 0.36, and 0.37 for the metacognitive module, Top2Diff, and SoftMax, respectively; Figure 6C). Thus, the metacognitive module not only allowed confidence to correct for bias but also made confidence ratings much more informative of the accuracy of the primary decision.

## Discussion

We tested whether human confidence judgments are bias-aware or bias-blind. To examine this, we used simulations to identify a signature of bias-aware confidence: a negative influence of response bias on confidence when controlling for accuracy. Across two multi-alternative classification conditions (4-choice and 8-choice), we observed that human confidence robustly exhibited this signature of bias-aware confidence. Additional analyses demonstrated that these results cannot be explained by guessing and that the strength of bias correction was attenuated under speed focus instructions that resulted in shorter confidence response times. In contrast, standard ANNs, which give confidence through mechanisms that do not correct for bias, exhibited bias-blind confidence. Importantly, augmenting the ANNs with a metacognitive module that learns to predict their accuracy and thus can correct for response bias allowed them to produce the signature of bias-aware confidence. Our results show that humans exhibit bias-aware confidence and that even though standard ANNs give confidence in a bias-blind manner, a simple metacognitive module allows ANNs to make confidence judgments that are correct for response bias.

### Implications for human metacognition

Traditional theories of human metacognition have emphasized the monitoring not only of decision uncertainty, but also of one’s own cognitive processes (Flavell, 1979; Nelson & Narens, 1994.; Shimamura, 2000). However, these self-monitoring processes have been studied mostly within the context of education and memory – fields that lack tight control over stimuli and thus are limited in the precision of computational models they can support. More recently, perceptual metacognition research has gained prominence due to its ability to build precise computational models using well-controlled stimuli (Rahnev, 2021). However, virtually all prominent models in visual metacognition only feature monitoring of stimulus uncertainty, with no role for monitoring of one’s own response bias (Rausch et al., 2023; Shekhar & Rahnev, 2024a). It has thus remained unclear whether human confidence is bias-aware or bias-blind in the context of controlled perceptual tasks.

Our work demonstrates that this gap is largely due to the fact that the field has almost exclusively focused on 2-choice tasks. Such tasks do not allow the estimation of a separate response bias associated with each choice, which then can be linked to confidence for that choice. Here, we employed 4- and 8-choice tasks that make it possible to compute category- specific response bias and confidence. Our results demonstrate that a clear signature of bias- aware confidence – a negative direct effect of response bias on confidence – emerges for human participants across multiple conditions and experiments.

Our results further showed that the degree of bias correction in confidence judgments was diminished under speed focus. The effect was presumably driven by the fact that the speed focus resulted in faster confidence response times (cRTs), which meant that confidence judgments had less time to perform bias-correcting computations that require the retrieval of estimated response biases and correcting for them in the final confidence rating. Given that confidence judgments are known to rely on the prefrontal cortex even for perceptual tasks (Middlebrooks & Sommer, 2012; Morales et al., 2018; Qiu et al., 2018; Yeon et al., 2020), it is plausible to assume that this bias correction is dependent on a limited cognitive resource and thus decreasing the time available for confidence judgments would decrease the degree of bias correction. In contrast, stimulus uncertainty information is already present in early visual cortex (Bergen & Jehee, 2019), supporting fast and largely automatic extraction of sensory uncertainty, which can then be used for confidence (Aguilar-Lleyda et al., 2021; Xue et al., 2023). Our findings thus open new ways of empirically dissociating different types of uncertainty signals (Fleming, 2021) even in simple perceptual tasks.

### Implications for ANN metacognition

Our results showed that standard ANNs trained on object recognition exhibit bias-blind confidence. The absence of bias correction was seen across ANN architectures (AlexNet, ResNet18, and VGG19) and despite matching their overall performance to that of humans. Indeed, this lack of bias monitoring in standard ANNs is not surprising: these networks derive confidence from instantaneous evidence on each trial, without considering their decision tendencies or response history. This result demonstrates that, although ANNs have proven adept at capturing many aspects of human confidence judgments (Green et al., 2026; Shekhar et al., 2025; Shekhar & Rahnev, 2024b; Webb et al., 2023), these similarities likely stem from both systems having to solve a common problem constrained by the same stimuli and tasks. However, to the extent that human confidence is based on cognitive mechanisms that go beyond the processing of the current stimulus, ANN and human confidence begin to diverge (Xue et al., 2026). Therefore, building human-like metacognition into neural networks requires that we go beyond emergent properties and reproduce the bias monitoring and other metacognitive mechanisms used by humans that go beyond processing stimulus uncertainty.

Several previous papers have built metacognitive capability in ANNs that goes beyond the direct readout of stimulus and decision uncertainty (T. Chen et al., 2019; Corbière et al., 2019). Our metacognitive module follows this broad approach by training a separate confidence readout to predict whether a frozen classifier’s response is correct. This model provides a computational test of whether a dedicated reliability-monitoring readout can recover the human-like signature of bias-aware confidence. Critically, because the base classifier was frozen, the metacognitive module altered only the confidence assigned to each response, not the perceptual decision itself. The metacognitive module allowed the ANNs to exhibit the signature of bias-aware confidence, matching the qualitative pattern observed in humans. Supplementary analyses showed that our specific metacognitive module is not unique and that other possible modules – such as a module trained on the logits in the penultimate instead of the last layer of each network – can also produce bias-aware confidence (Supplementary Figure 7). Thus, human-like bias-aware confidence can be recovered by adding dedicated metacognitive mechanisms, even though it does not emerge spontaneously from standard ANN confidence readouts.

## Conclusion

By employing multi-alternative task designs, our work reveals that humans produce bias-aware confidence that corrects for biased decision tendencies even in simple perceptual tasks. In contrast, standard ANNs give confidence based solely on stimulus and decision uncertainty. Nevertheless, human-like bias monitoring can be incorporated into ANNs via a metacognitive module. These results show that human perceptual metacognition engages complex bias monitoring mechanisms and paves the way for replicating these mechanisms in neural networks.

## Methods

### Human experiments

#### Experiment 1

For Experiment 1, we recruited adult participants online using Georgia Tech’s SONA system and the Prolific platform (https://www.prolific.com). Participants on SONA (n = 9) received 2 research credits for their time in the 2-day study. Participants on Prolific (n = 194) were compensated $12/hour. A total of 203 participants completed the study. Three participants were excluded because of using only one number on the confidence scale (e.g., only using the number 2 for all responses on the 1-4 scale). Therefore, a total of 200 participants (45.5% females, mean age: 43.5 years, SD: 13.8 years) were included in the analysis. The study was approved by the Georgia Tech Institutional Review Board, and all participants were provided written informed consent.

Participants performed a noisy digit recognition task in which grayscale digit images from the MNIST dataset (Deng, 2012) were corrupted with uniform noise (Figure 1A). The strength of the original image was controlled by a signal-weight parameter λ, such that the intensity of each pixel was given by: *I_final_* = λ × *I_original_* + (1 − λ) × *I_random_*, where *I_final_* is the final intensity of the pixel, *I_original_* is the original intensity of the pixel, and *I_random_* is the random value ranging from 0 to 255. Thus, larger values of λ preserved more of the original image and produced easier stimuli, whereas smaller values of λ introduced more visual noise and produced more difficult stimuli.

Visual noise was adjusted separately for each choice-set condition using a modified 2-up/1- down adaptive staircase procedure (Leek, 2001; Levitt, 1971). To compensate for differences in choice-set size, the staircase started from different λ values across conditions: λ = 0.75 in the 4- choice condition, and λ = 0.70 in the 8-choice condition. In both conditions, the staircase step size was 0.10 for the first 10 trials, 0.05 for the next 10 trials, and 0.01 for the remaining trials. The procedure tracked the number of consecutive correct responses. After the first correct response in a run, the stimulus difficulty was unchanged. After the second consecutive correct response, visual noise was increased by decreasing λ, making the stimulus more difficult. Importantly, unlike the standard up-down staircase, the consecutive-correct counter was not reset after this increase. Therefore, each additional correct response within the same correct run produced another increase in visual noise. In contrast, an incorrect response decreased visual noise by increasing λ and reset the consecutive-correct counter to zero. This modified rule targets approximately 62% correct and thus differs from the classic 2-down/1-up staircase, which targets approximately 71% correct.

On each trial, a fixation cross was presented for 700 ms, followed by a digit stimulus (300 ms). Participants then made an untimed perceptual decision indicating which digit had been presented. After the perceptual response, participants rated their confidence in the accuracy of their decision on a 4-point scale (1 = low confidence, 4 = high confidence). There were three conditions that varied in the number of categories. Specifically, we included 2-, 4-, and 8-choice conditions where participants discriminated between digits 5 and 6, digits 5–8, or digits 1–8, respectively. Note that only the 4- and 8-choice conditions were analyzed here.

Participants completed two separate days of testing. On each day, participants completed all three conditions twice in an ABCABC block design, where A, B, and C were randomly assigned to the three choice conditions for each participant. Each block contained 120 trials in the first cycle and 100 trials in the second cycle, yielding 220 trials per condition. To allow performance to stabilize, the first 20 trials of each condition were excluded from analysis, resulting in 200 analyzed trials per condition (600 trials total in each day). This left 200 analyzed trials per condition per day and 400 analyzed trials per condition across the two testing days. Thus, each participant contributed up to 400 trials to each of the 4- and 8-choice analyses. Untimed breaks were provided between blocks. The experiment was implemented using jsPsych (de Leeuw, 2015).

#### Experiment 2

To examine whether task demands modulate the relationship between response bias and confidence, we analyzed an additional, previously published experiment (Rafiei et al., 2024). All experimental details are available in the original publication; here we give a brief overview of the critical components of the experiment. Participants completed an 8-choice digit classification task that was very similar to the 8-choice condition in Experiment 1. On each trial, participants first fixated on a fixation cross, followed by the presentation of a handwritten digit embedded in visual noise for 300 ms. Participants then reported which digit (1–8) was presented and subsequently rated their confidence on a 4-point scale (1 = lowest confidence, 4 = highest confidence). Responses and confidence ratings were self-paced. Critically, Experiment 2 used a within-participant speed-accuracy trade-off manipulation, such that, in different conditions, participants were instructed to emphasize either accuracy or speed. The experiment included N=60 participants, each completing a total of 480 trials in the accuracy and 480 trials in the speed condition.

### Statistical analysis

Behavioral measures were summarized separately for each participant, experimental condition, and digit category. We quantified response bias as the false alarm rate (FAR). For a given digit category, FAR was defined as the proportion of trials on which that digit was selected when it was not the correct label. Higher FAR values therefore indicate a stronger tendency to select that category in its absence. Response bias showed substantial variability across observers and digit categories, with individual participants exhibiting different magnitudes and patterns of category-specific response tendencies (Supplementary Figure 8).

#### Category-level mixed effect regression model

To test whether response bias predicts confidence independently of perceptual performance and RT, we fit a series of linear mixed-effects models at the digit level. For each participant, condition, and digit category, we computed FAR, accuracy, RT, and confidence, and z-scored all variables prior to analysis. We then estimated three models predicting confidence using only response bias as a predictor (*Confidence* ∼ *FAR*), using both response bias and accuracy as predictors (*Confidence* ∼ *FAR* + *Accuracy*), and using response bias, accuracy, and RT as predictors (*Confidence* ∼ *FAR* + *Accuracy* + *RT*).

All models included participant as a grouping factor with random intercepts and random slopes for included predictors, allowing the estimation of individual differences in baseline confidence and predictor effects. Our primary quantity of interest was the regression coefficient for FAR, assessed across the three models to evaluate whether the bias-confidence relationship persisted after controlling for accuracy and RT. Analyses were performed in Python using the *statsmodels* package (Seabold & Perktold, 2010). We additionally assessed multicollinearity among regression predictors using variance inflation factors (VIFs). Across all conditions in both experiments, VIF values for FAR, accuracy, and RT ranged from 1.01 to 1.92 (Supplementary Table 1), well below commonly used thresholds for problematic multicollinearity. These results indicate that FAR, accuracy, and RT provided sufficiently independent predictors in the mixed- effects regression analyses.

#### Bayesian multilevel mediation analysis

To test whether the relationship between response bias and confidence operates indirectly through accuracy, we conducted a hierarchical Bayesian mediation analysis with accuracy as the mediator of the bias-confidence association. This analysis allowed us to estimate the relationship between response bias and accuracy (*a*), the relationship between accuracy and confidence (*b*), and, critically, the direct relationship between response bias and confidence after accounting for accuracy (*c*′). We used default priors. Specifically, we used *N* (0,1.5) for population-level intercepts, *N* (0,1) for population-level path coefficients, half-normal distributions with a scale parameter of 1 for the residual and random-effect standard deviations, and a Lewandowski-Kurowicka-Joe (LKJ) prior with shape parameter *η* = 2 for the random-effect correlation matrix.

Models were implemented in Python using the PyMC package (Abril-Pla et al., 2023). Posterior sampling used four Markov chain Monte Carlo (MCMC), with 1,500 tuning iterations followed by 1,000 retained draws per chain, producing 4,000 posterior draws. The target acceptance rate was 0.99. Convergence was evaluated using rank-normalized *R̂*, bulk and tail effective sample sizes, divergent transitions. We report posterior means and 95% Highest Density Intervals (HDIs) for the direct, indirect, and total effects.

#### Moderated regression and mediation analyses for accuracy- and speed-focused conditions in Experiment 2

Experiment 2 featured accuracy- and speed-focus conditions. Since our goal was to understand the effect of response bias on confidence under speed pressure, we analyzed the two conditions jointly rather than using independent models. This was done for both the regression and mediation analyses.

For the mixed-effects regression analysis, we fit models predicting confidence where each predictor included interaction with condition. The FAR-by-condition interaction tested whether the relationship between response bias and confidence differed between the instruction conditions. Condition-specific simple slopes were estimated from main effect by using the main effect and adding or subtracting the condition difference.

For the Bayesian mediation analysis, we included condition main effects and interactions between condition and each mediation path. These models produced condition-specific *a*, *b*, and *c*^′^ paths. The moderation of the direct effect was quantified as the difference between the accuracy- and speed-focus *c*^′^ paths. The index of moderated mediation was defined as the corresponding difference between the covariance-adjusted indirect effects. The random intercepts and random *a*, *b*, and *c*^′^ slopes followed the same correlated multilevel structure and priors used in the unmoderated mediation models.

### Multi-alternative signal detection theory (SDT) simulations

We used multi-alternative SDT simulations to examine the relationship between response bias and confidence. Simulations were conducted separately for *K* = 4 and *K* = 8 response alternatives. For each condition, we simulated 200 observers completing 400 trials each, matching the design of Experiment 1. On each trial *t*, the true category *k_t_* was sampled randomly from the *K* response alternatives. The evidence for the true alternative was sampled from a normal distribution *N* (*μ*,1), whereas evidence for each incorrect alternative was sampled from *N* (0,1). The parameter *μ* was calibrated separately for each choice-set size to produce accuracy that matches the Experiment 1 empirical accuracy (4-choice: 0.63; 8-choice: 0.63).

Each observer (*s*) was assigned a fixed, mean-centered response bias vector *b_s_* = (*b_s_*_,1_,…,*b_s_*_,*K*_) The *b_s_*_,*i*_ values were sampled from a normal distribution 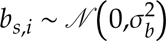 with standard deviation *σ_b_* and then centered so that ∑*_i_ b _s_*_,*i*_ = 0. Choices were generated by adding this response-bias vector, *b_s_*_,*i*_, to the evidence for each alternative, *x_s_*_,*t*,*i*_, and selecting the option with the largest biased decision variable: 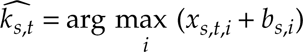. The magnitude of response bias, *σ_b_*, was calibrated separately for the 4- and 8-choice conditions so that the simulations reflect the overall level of bias in the behavioral data. Specifically, we matched the average participant- level standard deviation (SD) of FAR values for each condition, which were .070 for the 4-choice condition and .043 for the 8-choice condition. We used a bounded root search to identify the value of *σ_b_* that reproduced each empirical target. The resulting values were *σ_b_* = .368 for the 4- choice condition and *σ_b_* = .446 for the 8-choice condition. These values were fixed in all subsequent simulations.

The observers’ choices were determined by the biased decision variables (*x_s_*_,*t*,*i*_ + *b_s_*_,*i*_) and therefore did not change with the confidence model. To manipulate the degree of bias correction in confidence, we defined the evidence available to the confidence readout as *y_s_*_,*t*,*i*_ (*α*) = *x_s_*_,*t*,*i*_ + *b_s_*_,*i*_ − *αb_s_*_,*i*_ = *x_s_*_,*t*,*i*_ + (1 − *α*)*b_s_*_,*i*_, where *α* is the strength of bias correction with *α* = 0 indicating no bias correction and *α* = 1 indicating perfect bias correction. We varied *α* from 0 to 1 in increments of .1. Confidence was quantified using the Top-2 difference computation (Top2Diff), which has been shown to most closely match human confidence in multi-alternative tasks (Shekhar et al., 2025; Xue et al., 2024). Top2Diff simply takes the difference between the evidence for the chosen alternative evidence and the evidence for the strongest nonchosen alternative: 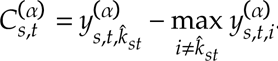 Note that bias correction could change the identity of the strongest competitor but never the observer’s original choice. In some cases, bias correction resulted in negative Top2Diff confidence values, indicating that the chosen alternative had lower corrected evidence than an unchosen alternative. Because human confidence ratings were given on a 4-point scale, such cases would correspond to the lowest confidence response category (confidence = 1). Therefore, negative simulated confidence values were interpreted as equivalent to the minimum confidence rating when comparing simulation results with human behavior.

The procedure above resulted in separate simulated datasets that matched the characteristics of the 4-choice and 8-choice conditions in Experiment 1 but with different levels of bias correction (*α* between 0 and 1). For each simulated dataset, we performed the mixed-effects regression and multilevel mediation exactly as done for the human data. We additionally quantified metacognitive sensitivity as the within-observer point-biserial correlation between trial-level confidence and correctness (Phi; Kornell et al., 2007; Rahnev, 2025). Metacognitive sensitivity was calculated separately for every simulated observer, choice-set size, and correction strength. Population means and 95% confidence intervals were estimated using 2,000 bootstrap resamples of the simulated observers.

### ANN modeling

#### Base ANN architectures and training

To examine whether standard artificial neural networks (ANNs) exhibit the same relationship between response bias and confidence as humans, we trained three convolutional architectures: AlexNet (Krizhevsky et al., 2012), ResNet18 (He et al., 2015), and VGG19 (Simonyan & Zisserman, 2015) to perform digit recognition using the MNIST dataset. These architectures were selected to represent a range of depth and structural complexity. All models were implemented in PyTorch (Paszke et al., 2019).

Because the MNIST digits are natively 28 × 28, all images were upsampled to 224 × 224 pixels using bilinear interpolation. The first convolutional layer of each network was modified to accept single-channel (grayscale) input, and the final classification head was replaced with a linear layer outputting 10 unnormalized logits (corresponding to digits 0-9).

We trained N = 60 independent instances of each architecture to support between-subject statistical analyses. Models were trained on the standard MNIST training set using the AdamW optimizer (Loshchilov & Hutter, 2019) with learning rate = 1×10⁻⁴ and weight decay = 1×10⁻⁴. To prevent overfitting and ensure optimal convergence, we employed a dynamic learning rate scheduler that halved the learning rate whenever validation loss plateaued for 2 consecutive epochs. Data augmentation was applied during training to encourage robust feature learning, including random rotations (±10°) and translations (up to 10% of image size). Training persisted for up to 20 epochs with early stopping (patience = 5 epochs), and the model state with the highest validation accuracy was retained for analysis. Batch sizes were set to 128 for AlexNet and ResNet18, and 64 for VGG19 to accommodate memory constraints; this variation was purely computational and did not alter the optimization objective.

#### Psychophysical evaluation and confidence readout

Models were evaluated on the full MNIST test set (10-way classification). To parallel the human experiment, Gaussian noise was added to test images. For each architecture, we identified the noise level that yielded overall classification accuracy comparable to human performance in Experiment 1.

We used both the Top2Diff and the SoftMax methods to derive ANN confidence. Specifically, for each image *i*, the base classifier produced a vector of logits ***l****_i_*. These logits were converted into class probabilities using the SoftMax function: 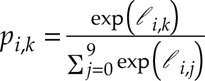. The network’s response was the category with the largest logit: 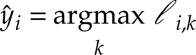 The Top2Diff confidence was based on the difference between the two largest logit values: 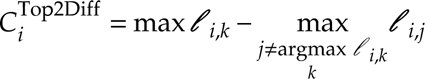

The SoftMax confidence was based on the maximum SoftMax probability: 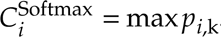 . Thus, the two standard confidence readouts (Top2Diff and the SoftMax) were both derived directly from the base classifier’s logits and did not involve separately learned confidence mechanisms. We treated each ANN instance as a separate observer and performed the mixed- effects regression and multilevel mediation exactly as done for the human data. This was done separately for each model architecture and confidence readout: Top2Diff, SoftMax, and metacognitive module (see next section). Finally, we also computed the metacognitive sensitivity (Phi) associated with each of the three types of confidence readout.

#### Metacognitive module

We next examined whether a separately trained metacognitive module could produce the bias- aware confidence signature observed in humans. For each ANN instance, all classifier parameters were frozen, and the ANN was maintained in evaluation mode throughout metacognitive training. Consequently, training the metacognitive module could not change the classifier’s representations, logits, choices, accuracy, or response tendencies.

For each noisy training image, the frozen classifier produced logits ***l****_i_* and the metacognitive module used the ten logits to predict if the classifier’s response would be correct or not. The module consisted of a fully connected layer with 512 hidden units, a rectified linear activation, and a sigmoid output unit: *Ĉ_i_* = *σ* (W_2_ ReLU (W_1_ l*_i_* + b_1_) + *b*_2_) . The output *Ĉ_i_* represented the predicted probability that the frozen classifier’s response was correct. Because the module received the full logit vector rather than only the winning value or output margin, it could learn category-specific relationships between the classifier’s output patterns and its probability of being correct. The module was trained by minimizing binary cross-entropy between predicted confidence and response correctness, together with a weak regularization term discouraging degenerate confidence estimates.

Metacognitive training used noisy images from a class-balanced subset of the MNIST training split. Ten percent of these images were held out for validation. The same architecture-specific noise levels used during final evaluation were applied during metacognitive training. Only the parameters of the metacognitive head were optimized, using Adam with a learning rate of 1×10⁻⁴ and weight decay of 1 × 10⁻³. Training continued for 20 epochs, with a learning-rate reduction factor of 0.5 after three epochs without improvement in validation loss. Batch sizes were 128 for AlexNet and ResNet18 and 64 for VGG19. The final saved metacognitive head was used for evaluation.

## Data and code

Data and code are available at https://osf.io/nz25w/overview?view_only=36e5bcc2225f4b55a54b77b5f690d786 and https://github.com/bogeng-song/RespBias-Metacognition.

## Author contributions

B.S. designed the study, performed analyses, and wrote the manuscript.

D.R. supervised the project and revised the manuscript.

## Acknowledgments

This work was supported by the National Institutes of Health (award: R01MH119189). We thank Rachel Denison for helpful discussions about these results.

