## Supplemental figures for "Bias-aware versus bias-blind confidence in humans and machines"

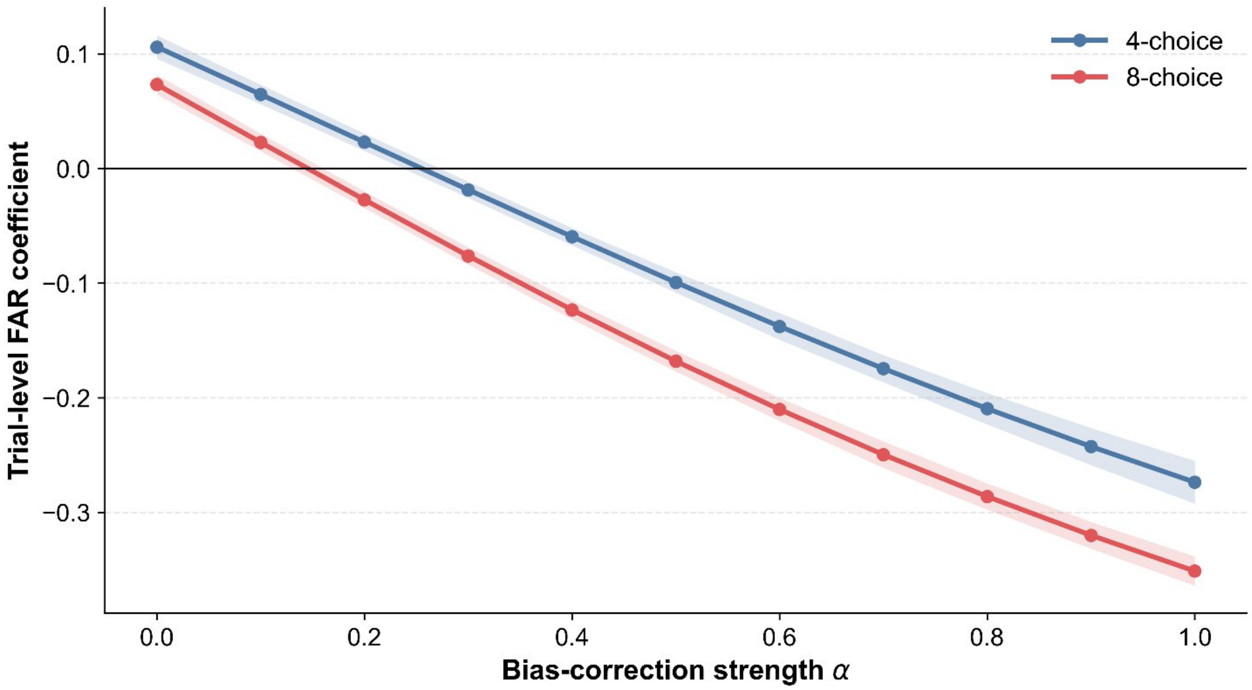


**Supplementary Figure 1. Trial-level response bias regression reveals an alternative signature of bias-aware confidence in simulations.** Simulated observers performed 4- and 8-choice tasks across varying bias-correction strengths ($\alpha$). Instead of running regressions on the data averaged across digits, here we explored an alternative trial-level analysis. Because there is no trial-by-trial measure that corresponds to response bias, each trial was assigned the false alarm rate (FAR) associated with the alternative selected on that trial. Trial-level FAR and confidence were standardized within each simulated observer. At each choice-set size and bias-correction strength, confidence was predicted from chosen alternative FAR using a linear mixed-effects model with observer-specific random intercepts and random slopes for FAR. The plotted values represent the fixed-effect FAR coefficients. Under bias-blind confidence ($\alpha$ with a small value), alternatives with higher FARs were associated with higher confidence. As bias correction ($\alpha$) increased, the FAR coefficient progressively decreased and became negative, indicating that confidence was increasingly discounted for alternatives associated with stronger response tendencies. Similar patterns were observed in the 4- and 8-choice simulations. Note that this trial-level analysis produces negative FAR coefficients at lower levels of bias correction (i.e., at smaller $\alpha$ values compared to our main analyses; Figure 2), which makes this test more sensitive but also increases the chance that bias-blind confidence may not exhibit significantly positive FAR coefficients due to noise. Shaded regions indicate 95% confidence intervals.

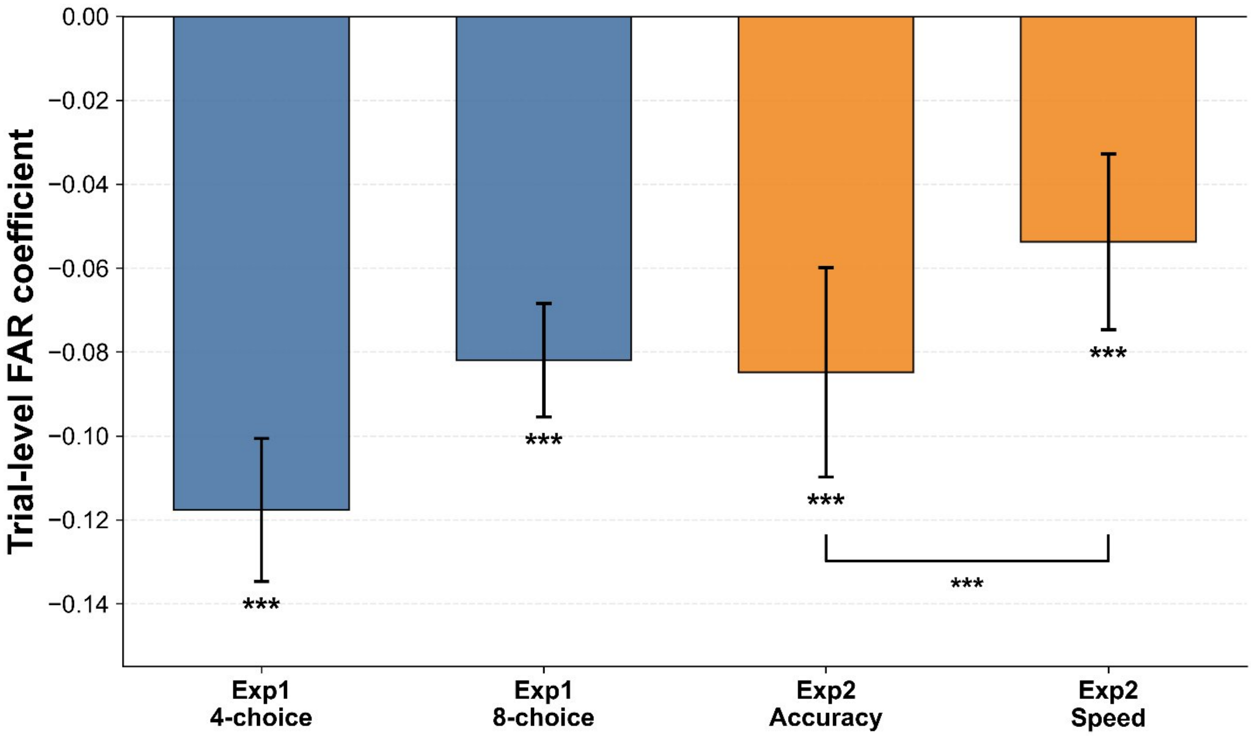


**Supplementary Figure 2. Trial-level response bias predicts lower confidence in humans.** We performed the alternative trial-level regression from Supplementary Figure 1 on the data from Experiments 1 and 2. This analysis was run exactly as in the main analyses from Figure 3B. Specifically, for Experiment 1, confidence was predicted from chosen alternative FAR using separate linear mixed-effects models for the 4- and 8-choice conditions. For Experiment 2, the accuracy- and speed-focus conditions were analyzed jointly using a FAR-by-condition interaction. Instruction condition was effect-coded as +0.5 for accuracy focus and −0.5 for speed focus, such that the interaction coefficient represented the accuracy-focus FAR slope minus the speed-focus FAR slope. All models included participant-specific random intercepts and random slopes for FAR, with no additional predictors. We found that FAR negatively predicted confidence in both conditions of Experiment 1 and under both instruction conditions in Experiment 2, indicating that participants exhibited the signature of bias-aware confidence in all four conditions. Note that our main analysis showed a FAR coefficient close to zero for the speed condition of Experiment 2, which we argued represents bias-aware confidence with smaller bias correction. The fact that this more sensitive analysis produces a negative FAR coefficient further supports that conclusion. Finally, we also confirmed that the negative FAR coefficient was significantly different under accuracy focus than under speed focus (FAR × Condition: $\beta = -0.031$, $SE = 0.008$, $z = -3.75$, $p < .001$, 95% CI [−0.047, −0.015]), again showing that speed focus reduces the strength of bias correction. Bars show the estimated fixed-effect FAR slopes, error bars indicate 95% confidence intervals, asterisks beneath individual bars denote simple-slope tests against zero, and the bracket denotes the FAR-by-condition interaction. ***, $p < .001$.

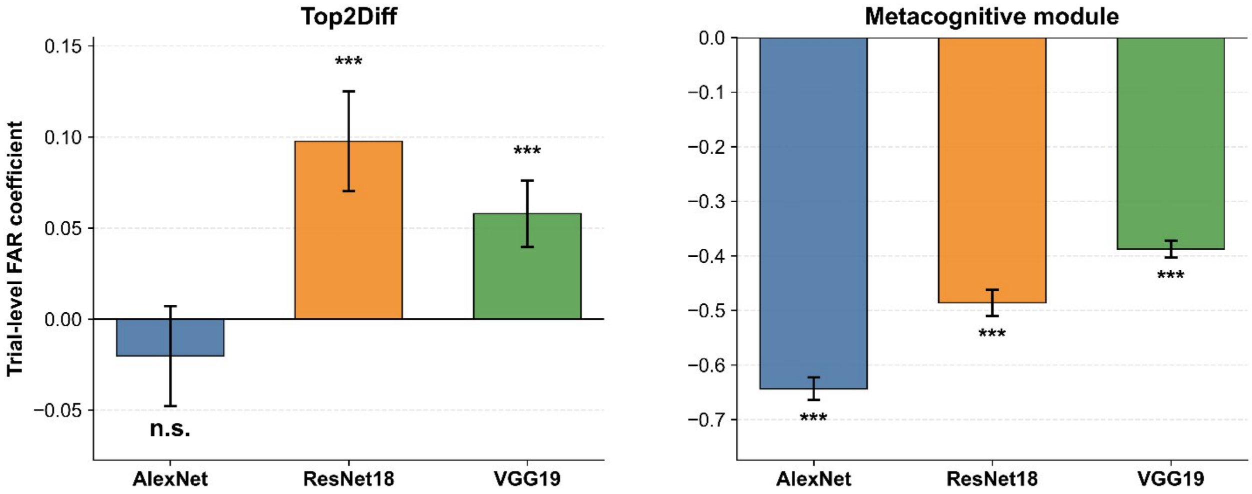


**Supplementary Figure 3. Trial-level response bias reveals bias-blind and bias-aware confidence readouts in ANNs.** We performed the alternative trial-level regression from Supplementary Figure 1 on the data from the three ANN architectures. Separate linear mixed-effects models were fitted for each architecture and confidence readout, with confidence predicted from chosen-alternative FAR and with instance-specific random intercepts and random slopes for FAR. Bars show the estimated fixed-effect FAR coefficients. Left: For Top2Diff confidence, defined as the difference between the two largest logits. FAR positively predicted confidence in ResNet18 and VGG19, consistent with a bias-blind readout. Note, however, that this effect was not significant for AlexNet, which may be due to the fact that the trial-level analysis is more likely to fail to show a positive FAR coefficient for bias-blind confidence (Supplementary Figure 1). Right: For confidence produced by the metacognitive module, FAR negatively predicted confidence across all three architectures, indicating that the module produced bias-aware confidence judgments. Error bars indicate 95% confidence intervals. ***, $p < .001$; n.s., not significant.

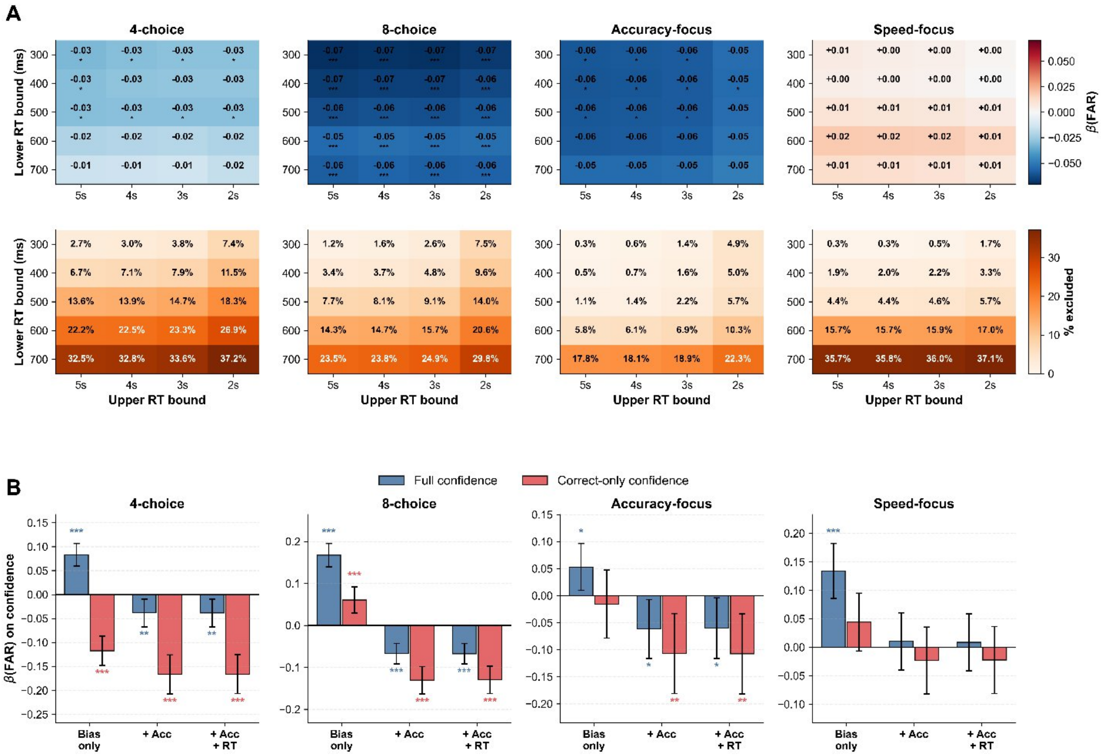


**Supplementary Figure 4. Control analyses to test whether the main effects were driven by guessing.** Our main results demonstrated that response bias negatively predicted confidence when accuracy was controlled for in both Experiments 1 and 2. We argued that this pattern of results reflects a bias-aware confidence strategy. However, a possible alternative explanation of these results is that they are due to biased guessing accompanied by low confidence. Specifically, a strategy where participants always choose the same default digit when uncertain and report low confidence on such trials could also produce a negative relationship between response bias and confidence. Here we perform two additional analyses that suggest that our results were not driven by such a strategy. (A) Guesses are typically accompanied by either very fast or very slow responses (Ratcliff, 1993; Ratcliff & Tuerlinckx, 2002; Unsworth et al., 2010; Yellott, 1971).Therefore, we repeated the main regression after applying exclusion criteria for fast RTs (300, 400, 500, 600, and 700 ms) and slow RTs (2, 3, 4, and 5 seconds). We found that the negative FAR confidence relationship pattern remained stable across RT windows for all conditions. Thus, the signature of bias-aware confidence remained even after excluding the trials that are most likely to reflect guessing. The top row shows the FAR coefficient after controlling for accuracy and RT for each RT window. The bottom row shows the percentage of trials excluded by each RT window. (B) Correct only trials confidence analysis. The original regressions were repeated after computing confidence using only correct trials, while FAR, accuracy, and RT were computed as in the original analysis. Note that guesses in multi-alternative tasks typically result in errors and thus would have minimal influence on averages based on correct trials only. The results replicated our findings when all trials were used: we found negative FAR-confidence relationship after controlling for accuracy, and after controlling for accuracy and RT, for both conditions in Experiment 1 and for the accuracy-focus conditions in Experiment 2, but no significant effect for the speed-focus condition in Experiment 2. Error bars indicate 95% confidence interval. *, p < .05; **, p < .01; ***, p < .001.

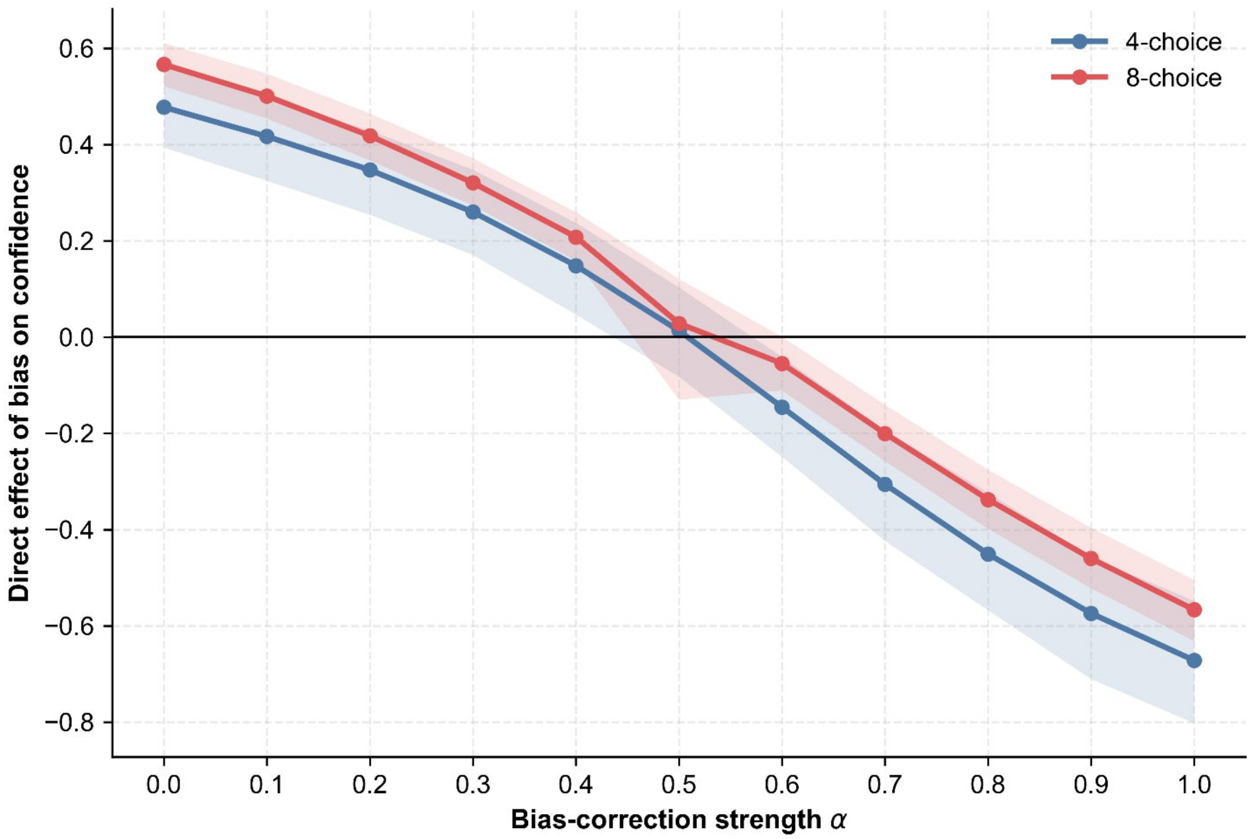


**Supplementary Figure 5. Bias correction strength determines the direct effect of response bias on confidence in simulations.** Bayesian mediation analyses were used to examine how the direct relationship between response bias and confidence changes as a function of bias-correction strength ($\alpha$). For each simulated observer, response bias influenced confidence through two pathways: an indirect pathway mediated by accuracy and a direct pathway representing the relationship between response bias and confidence after accounting for accuracy. This figure shows the estimated direct effect of response bias on confidence across levels of $\alpha$ for 4-choice and 8-choice simulations. At low bias-correction strength, the direct effect was positive, indicating that confidence increased for alternatives with stronger response tendencies even after accounting for accuracy. As bias correction increased, the direct effect progressively decreased and became negative, indicating that confidence was increasingly reduced for alternatives associated with stronger response tendencies. These results suggest that a direct effect of zero in the mediation analyses corresponds to a smaller bias correction instead of no bias correction and therefore the close-to-zero effect for the speed focus condition in Experiment 2 (Figure 4C) indicates reduced rather than eliminated the bias-correction within confidence judgments. Shaded regions represent 95% highest-density intervals.

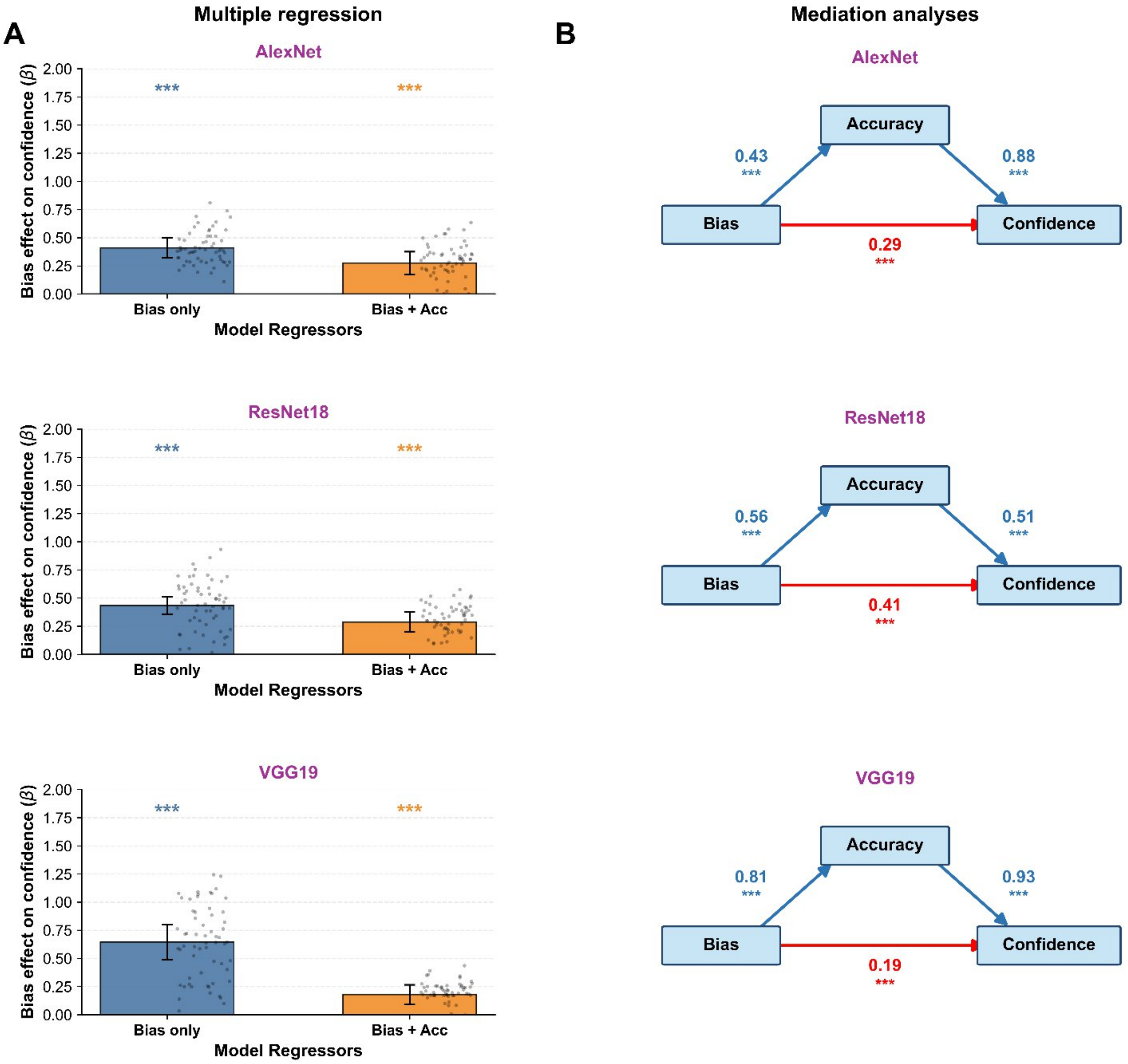


**Supplementary Figure 6. SoftMax confidence in standard ANNs replicate the Top2Diff results.** In our main paper, we quantified confidence in ANNs using the Top2Diff computation. Here we show that the results remain the same when instead using SoftMax as the confidence measure. For each ANN architecture, confidence was quantified as the maximum SoftMax probability assigned to the selected category. (A) Mixed-effects regression analyses testing whether response bias predicts confidence. The bias-only model estimates the overall association between FAR and confidence, whereas the bias + accuracy model tests whether response bias predicts confidence after accounting for differences in perceptual accuracy. AlexNet, ResNet18, and VGG19 exhibited the signature of bias-blind confidence. Individual points represent estimates from individual network instances; error bars indicate 95% confidence intervals; ***, $p < .001$. (B) Mediation analyses examining whether the relationship between response bias and confidence was mediated by accuracy. Path coefficients indicate posterior mean effects. Across all architectures, response bias increased accuracy and accuracy increased confidence, but a positive direct effect of bias on confidence remained, consistent with a bias-blind confidence mechanism, *** represent 99.9% highest-density intervals excluded zero.

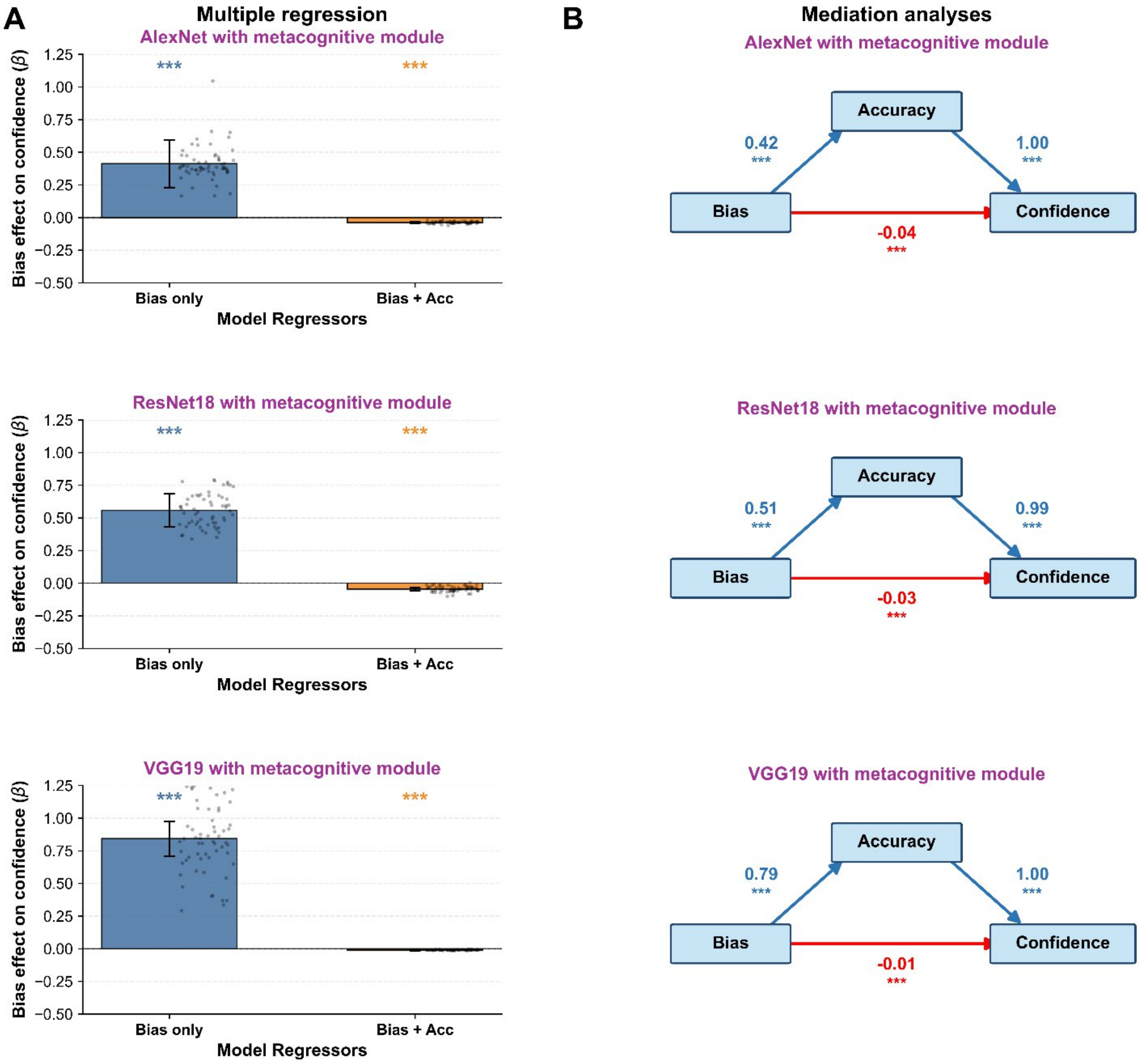


**Supplementary Figure 7. An alternative metacognitive module also produces bias-aware confidence in ANNs.** Our metacognitive module was trained to predict the accuracy of each classification based on the logits in the last layer. Here we show that an alternative 9 enitivenitive module that is based on the activations in the penultimate network layer also produces bias-aware confidence. For each AlexNet, ResNet18, and VGG19 instance, the parameters of the trained classifier were frozen, and a metacognitive head was trained to predict whether the classifier's response was correct using only its penultimate-layer representation. The input dimensionality was 4,096 for AlexNet and VGG19 and 512 for ResNet18. Dropout with a rate of 0.5 was applied to these representations before they were passed to the metacognitive head. The penultimate-only modules used the same correctness targets, noisy images, optimization procedure, and evaluation data as the metacognitive module from the main paper. Because the underlying classifiers remained frozen, metacognitive training did not alter their perceptual representations, classification responses, accuracy, or response biases. (A) Mixed-effects regression analyses tested whether categorylevel response bias predicted metacognitive confidence before and after accounting for accuracy. The results show that FAR positively predicted confidence in the bias-only models across architectures. After accuracy was included, the FAR coefficients became negative. Bars show the mean FAR slopes across network instances, points show instance-specific slopes, and error bars indicate 95% confidence intervals; ***, $p < .001$. (B) Bayesian multilevel mediation analyses separated the indirect association between response bias and confidence through accuracy from the direct association remaining after accuracy was accounted for. Across architecture, the direct paths from response bias to confidence were negative. Together with the primary metacognitive module results, these findings show that bias-aware confidence can be learned from different internal classifier representations without modifying the underlying perceptual decision process. Path coefficients are posterior means, and asterisks indicate that the corresponding 95% highest-density intervals excluded zero.

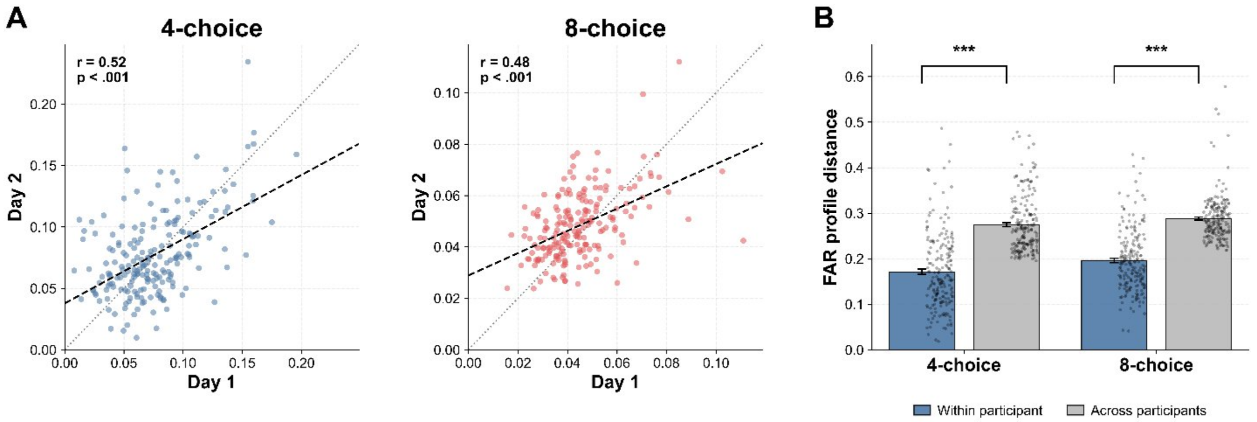


**Supplementary Figure 8. Individual differences in response bias are stable across days.** We tested for the existence of stable individual difference in response bias in two analyses, each comparing response bias across the two days of testing. (A) We first tested whether participants exhibited individual differences in the magnitude of response bias. For each participant, the overall magnitude of response bias was calculated separately for Day 1 and Day 2 as the standard deviation of the digit-specific FARs. Response-bias magnitude was positively correlated across days in both the 4-choice condition (Pearson's $r = .52$, $p < .001$) and the 8-choice condition ($r = .48$, $p < .001$), indicating that participants who exhibited greater variability in response tendencies on Day 1 tended to exhibit greater variability on Day 2. Black dashed lines show the fitted regression lines; gray dotted lines show the identity line; dots represent individual participants. (B) In a second set of analyses, we tested whether participants exhibited individual differences in the exact profile of digit-specific response bias. The stability of each participant's digit-specific response-bias profile was quantified as the sum of the absolute differences between corresponding FAR values across days. Within-participant distance compared each participant's Day 1 profile with their own Day 2 profile. Across-participant distance compared each participant's Day 1 profile with the Day 2 profile of every other participant; these distances were then averaged to produce a single across-participant average distance per participant. Each participant therefore contributed one within-participant and across-participant distance. Within-participant distances were significantly smaller than across-participant distances in both choice conditions, indicating that individual response-bias profiles were more similar to themselves across days than to the profiles of other participants. Bars show group means, points show participant-level distances, and error bars indicate SEM. ***, $p < .001$.

| Condition | N | VIF FAR | VIF Accuracy | VIF RT | Max VIF |
|---|---|---|---|---|---|
| Expt 1, 4 choices | 200 | 1.92 | 1.92 | 1.01 | 1.92 |
| Expt 1, 8 choices | 200 | 1.49 | 1.49 | 1.01 | 1.49 |
| Expt 2, accuracy focus | 60 | 1.05 | 1.04 | 1.01 | 1.05 |
| Expt 2, speed focus | 60 | 1.14 | 1.06 | 1.21 | 1.21 |

**Supplementary Table 1. Multicollinearity diagnostics for the main multiple-regression models.** Variance inflation factors (VIFs) were computed for the predictors in the full regression model predicting confidence from FAR, accuracy, and RT. Rows indicate the number of subjects × response-category observations included in each condition. The maximum VIF was 1.92, and all condition numbers were below 1.5, indicating low multicollinearity among predictors.
